# 3D Printed Molds for Organ-on-a-Chip and Fluidics: PDMS-Based Rapid and Accessible Prototyping

**DOI:** 10.1101/2025.03.29.645830

**Authors:** Rana J. Abbed, Edwin I. Quiñones Cruz, Susan E. Leggett

## Abstract

The ability to rapidly fabricate custom polydimethylsiloxane (PDMS) devices is central to advancing organ-on-a-chip (OoC) technologies and other biological microplatforms. However, traditional photolithography and the surface roughness of directly 3D printed molds limit their accessibility and scalability of PDMS-based systems. Photolithographic workflows are limited by their dependence on specialized equipment, technical expertise, dedicated fabrication infrastructure, and are typically restricted to planar geometries and microscale features, limiting their use for millifluidic or complex 3D device features. To address these challenges, we present a modular workflow for the robust fabrication of PDMS-based devices using stereolithography (SLA) or fused deposition modeling (FDM) printing combined with optimized epoxy coatings. Acetone-thinned epoxy formulations dramatically improve SLA printed mold smoothness, eliminate tearing during demolding, and yield PDMS replicas with clean, well-defined structural features. For FDM printed molds, a two-step epoxy coating strategy restores mold quality sufficient for robust replica molding. The resulting PDMS devices support irreversible glass bonding, fluid containment, and cell culture applications, validated using normal mammary epithelial and cancer cell lines. We further demonstrate the formation of perfusable tissue aggregates within 3D matrices and introduce a low-cost 3D printed imaging platform for parallel live-cell imaging across four PDMS devices, showcasing its use for monitoring 20 OoC channels under gravity- or pump-driven flow. This versatile and reproducible method lowers the barrier to entry for soft lithography, allowing researchers without prior microfabrication expertise to rapidly prototype functional PDMS devices for diverse biological applications.

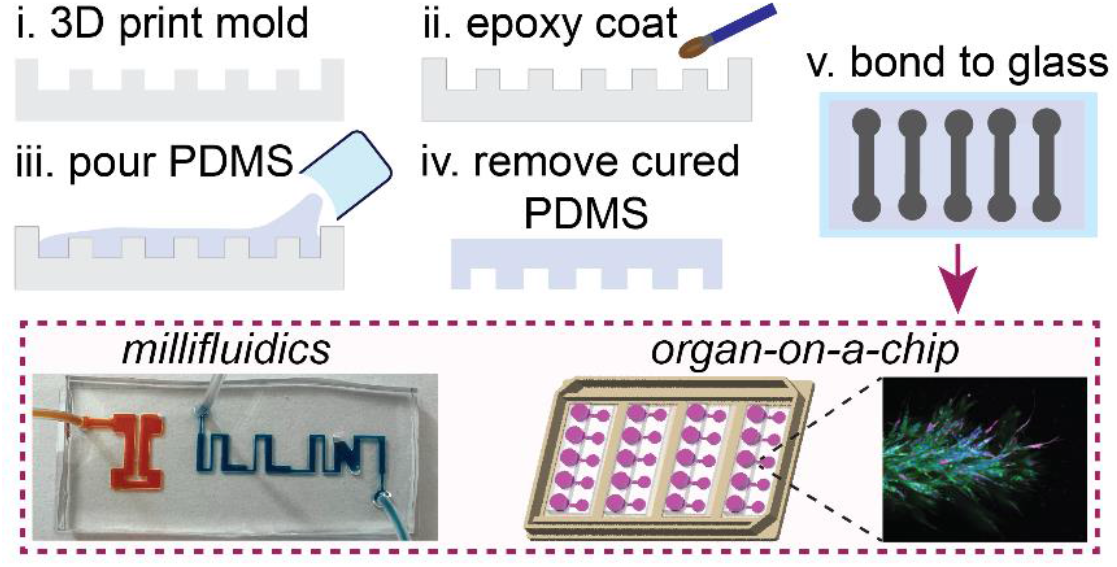

## Introduction

The fabrication of devices with complex three-dimensional (3D) structures has been critically enabled by rapidly advancing 3D printing technologies, and is particularly impactful in engineering, biomedical research, and medicine as a highly customizable and cost-effective approach for rapid prototyping.^1^ Among various 3D printing technologies, stereolithography (SLA) has emerged as a powerful tool for producing precise microscale structures, with typical layer thicknesses of commercial printers as fine as 25 microns.^2, 3^ It has fast-tracked the fabrication of microfluidic devices for applications such as biochemical assays, diagnostics, and droplet technologies,^4^ where precision and scalability are critical. Additionally, fused deposition modeling (FDM) provides a cost-effective, rapid, and facile approach to fabricating structures with meso-scale to millimeter-scale features. However, photolithography has persisted as the primary approach for producing molds for on-chip fluidic devices for cell culture applications, despite its several disadvantages compared to 3D printing – photolithography typically has higher fabrication costs, is more time-consuming, requires greater technical expertise, and is less amenable to rapid prototyping. Moreover, photolithography is generally limited to microscale feature sizes and is not robust for the generation of larger millimeter-scale structures essential for many biological applications, such as chambers and channels housing tissue biopsies, engineered tissues, and 3D cell cultures.

3D printing technology, including SLA and FDM, provides a cost-effective, rapid, and facile approach to fabricating molds for fluidic and organ-on-a-chip (OoC) devices.^5, 6^ Unlike traditional photolithography, which usually requires cleanroom facilities and extensive training, both SLA and FDM printing can be performed rapidly with minimal equipment in a standard laboratory environment, significantly reducing fabrication time and costs.^7, 8^ This accessibility makes them attractive alternatives for researchers optimizing new device designs and configurations without the constraints of traditional photolithography-based methods.^7, 9^ Recent studies have demonstrated the potential of 3D printing to address these limitations by enabling rapid production of complex geometries with a user-friendly workflow.^8, 10–15^ Nevertheless, existing challenges remain for the direct use of 3D printed devices in cell culture systems, including potential cytotoxicity and high material rigidity. For instance, SLA printed devices may lack the mechanical flexibility needed for dynamic fluid control, while FDM printed devices often suffer from poor optical transparency and difficulty in sealing.^16, 17^ Polydimethylsiloxane (PDMS) has served as a longstanding material for biological applications, exhibiting viscoelastic properties and biocompatibility, enabling the fabrication of flexible fluidic devices, tissue engineering scaffolds, and biointerfaces that mimic soft tissue mechanics.^18^

Accordingly, generating PDMS devices from 3D printed molds represents an ideal workflow supporting diverse cell culture applications. However, both SLA and FDM 3D printed molds have inherent limitations that affect the quality and performance of the final PDMS-based devices.^19, 20^ One primary challenge is achieving high feature fidelity, as the layer-by-layer printing process introduces surface roughness and stair-stepping effects, particularly on curved or angled features.^21–23^ These imperfections impact the precision of microchannels and other critical structures, potentially altering the cellular microenvironment in OoC devices.^24–27^ FDM printing, in particular, results in high surface roughness and visible layer lines unsuitable for applications requiring smooth channel walls. Additionally, the direct use of 3D printed parts as molds presents challenges for PDMS replica fabrication, including potential cytotoxicity and reduced optical transparency. These issues may arise from the inherent properties of resins used in SLA and common filaments like polylactic acid (PLA) in FDM, further limiting their suitability for biological applications.

To address these limitations, we established a 3D print post-processing method for the fabrication of PDMS-based devices from 3D printed molds, optimizing epoxy formulations and application methods for coating both SLA and FDM printed molds. We demonstrated that clean PDMS replicas can be generated from SLA printed molds coated with a single acetone-thinned epoxy coating, or from FDM printed molds with a two-step epoxy coating, which involved the application of a standard epoxy mixture followed by the application of an acetone-thinned epoxy mixture. We generated molds for fluidic channel sizes ranging from micro-to millifluidic scales, as well as arrays of varied geometric posts. Our optimized workflow supports robust and reproducible fabrication of fluidic devices with feature dimensions spanning millimeter-scale chambers down to microscale channels. Critically, our PDMS devices demonstrate biocompatibility, optical clarity, and mechanical robustness suitable for 2D and 3D cell culture, fluid flow, and microscopy, bridging a critical scale gap for diverse biological applications. Finally, we established an OoC platform that enables fabrication and live-cell imaging of 20 OoCs in parallel. All 3D printed components can be produced using a basic desktop printer, greatly reducing the barrier to OoC technologies by eliminating the need for specialized equipment or prior microfabrication experience for facile PDMS-based device fabrication.

Compared to traditional photolithography, our method significantly reduces fabrication time and costs while enabling rapid prototyping in standard laboratory environments. Furthermore, our approach ensures mold durability, allowing multiple reuse cycles without degradation, thus greatly enhancing cost-efficiency. Overall, rather than directly using 3D printed parts as molds, our approach leverages epoxy-coated SLA and FDM printed molds to create high-quality molds for casting PDMS, maintaining the advantages of using PDMS-based devices while effectively overcoming traditional photolithography limitations. The practical, detailed methods provided here further support adoption and replication, positioning our approach as a highly accessible and valuable resource within the microfluidics, millifluidics, and biomedical communities.

## Methods

### SLA and FDM 3D Printing

All 3D printed parts and mold designs were created using SOLIDWORKS and generated by SLA or FDM printing. The Form 3+ (Formlabs) was used for stereolithographic printing with Clear V4 resin. 3D printing parameters were set to 25 µm layer height for molds incorporating <1 mm feature sizes, or 100 µm layer height for molds with ≥1 mm minimum feature size. After printing, molds were carefully removed and washed in a Formlabs Wash Station with 100% isopropyl alcohol (IPA) for 15 minutes. Following this initial rinse, the molds were transferred to the Formlabs Form 3 Finishing Kit and curing station for an extended soak in 100% IPA, lasting 4–6 hours to ensure thorough cleaning and resin removal. After soaking, the SLA printed molds were air-dried on a benchtop overnight, aided by a fan to expedite evaporation of residual IPA. FDM printing was used as an alternative to SLA printing. FDM printed molds were generated using the UltiMaker S5 with PLA filament, specifying printing parameters of 200 µm layer height and 20% infill. Images of 3D printed molds and resulting PDMS replicas were imaged using an iPhone 13 Pro camera.

### Standard Epoxy Coating of 3D Printed Molds

Two distinct two component epoxy systems were used to coat 3D printed molds in this study: XTC-3D™ (Smooth-On, Inc.) and Crystal Clear Epoxy Resin Kit (JANCHUN). For conciseness, these systems are referred to as ‘XTC-3D’ and ‘Janchun’ throughout the text. According to the manufacturers, XTC-3D is a 2:1 resin-to-hardener ratio system (2 parts A:1 part B by volume), whereas Janchun is a 1:1 system (1 part A:1 part B by volume). Accordingly, standard epoxy coatings for 3D printed molds were prepared as a two-part mixture with components A and B mixed at a 2:1 ratio for XTC-3D, or as a two-part mixture of components A and B at a 1:1 ratio for Janchun. Epoxy mixtures were applied using a disposable flat synthetic nylon paintbrush to deposit a thin, even coating on the surface of 3D printed molds.

To achieve uniform coverage, the brush was initially loaded with epoxy, which was then spread finely across the surface; any excess epoxy was removed from the brush before further application. Molds were brushed in alternating, perpendicular directions three times to ensure an even layer of epoxy coating. To minimize excess epoxy buildup, a controlled stream of air from a laboratory gas line was applied through tubing with a 200 µL pipette tip at the end to precisely direct airflow across mold features, ensuring a smooth, thin finish. Epoxy coating and air smoothing were performed under a chemical fume hood. Next, the epoxy-coated molds were left to cure overnight at room temperature. Alternatively, the epoxy coating on 3D printed molds could be cured at elevated temperatures to significantly reduce drying time from 12 hours to 1–2 hours. Molds coated with Janchun were cured at 65 °C for 1–2 hours, while molds coated with XTC-3D epoxy were cured at 40 °C for 1–2 hours. The elevated temperatures used reflect the epoxy manufacturer’s recommendations for drying temperature range, using the midpoint temperature to avoid potential issues that may occur at maximum temperature, such as melting. Once the epoxy had completely cured, coated molds were allowed to cool at room temperature for 10 minutes before adding PDMS to produce replicas.

### Epoxy Thinning for 3D Printed Molds

To achieve varying epoxy coating thicknesses, acetone was added to epoxy at different ratios to thin the mixture. Two specific formulations were prepared for both epoxy types, referred to as thin and thinner epoxy coatings. To obtain a thin coating, 2 parts of Part A, 1 part of Part B, and 0.25 parts of acetone (2A:1B:0.25 acetone), were measured and mixed in a plastic cup for XTC-3D epoxy, while 1 part of Part A, 1 part of Part B, and 0.25 parts of acetone (1A:1B:0.25 acetone), were mixed for Janchun epoxy. Similarly, for the thinner coating, 2 parts of Part A, 1 part of Part B, and 0.50 parts of acetone (2A:1B:0.5 acetone), were used for XTC-3D epoxy, and 1 part of Part A, 1 part of Part B, and 0.50 parts of acetone (1A:1B:0.5 acetone), were used for Janchun epoxy. The epoxy-acetone mixtures were thoroughly stirred for 1–2 minutes until homogeneous. The prepared mixtures were applied to their respective molds using a paintbrush, ensuring a uniform and thin application without excess material, as previously described. Following epoxy application, the molds were then dried at room temperature or under elevated temperatures (**Table 1**). Epoxy curing status was first assessed through physical methods, including tactile inspection for a dry, tack-free surface and a mechanical resistance test using light thumb pressure to ensure no deformation or imprinting. Chemical stability was verified via a solvent resistance test using isopropanol, with no surface softening or smearing indicating full cure.

**Table 1.** Working and Drying Tinies of XTC-3D and Janchun Epoxy Mixtures with Varying Acetone Ratios.

| XTC-3D epoxy |  |  |  |
| --- | --- | --- | --- |
| Acetone Ratio (parts) | Working Time (minutes) | Drying Time (Room Temperature) | Drying Time (40 $^{\circ}\text{C}$) |
| 0 | ~4-6 | 6 hours | ~2.5 hours |
| 0.25 | ~5-6 | 8-10 hours | ~2-2.5 hours |
| 0.50 | ~6-7 | 10-12 hours | ~2 hours |
| Janchun Epoxy |  |  |  |
| Acetone Ratio (parts) | Working Time (minutes) | Drying Time (Room Temperature) | Drying Time (65 $^{\circ}\text{C}$) |
| 0 | ~4-6 | 6 hours | ~1.5 hours |
| 0.25 | ~5-6 | 8-10 hours | ~1-1.5 hours |
| 0.50 | ~6-7 | 10-12 hours | ~1 hours |

### Multilayer Epoxy Coating of FDM Printed Channels

FDM printed molds were coated with either no epoxy, one layer of epoxy, or two layers of epoxy. Coating conditions were divided into five groups: (1) no epoxy control, (2) one layer of epoxy only, (3) one layer of epoxy 0.5 acetone mixture, (4) two sequential epoxy coatings, with epoxy only as the first layer and epoxy 0.5 acetone mixture as the second layer, and (5) two sequential epoxy coatings, with epoxy 0.5 acetone mixture as the first layer and epoxy 0.5 acetone mixture as the second layer. For conditions where two sequential epoxy coatings were applied, each layer of coating was allowed to fully cure before the application of the second coating, following curing times in **Table 1**. These epoxy coating conditions (groups 2-5) were performed for both XTC-3D epoxy and Janchun epoxy.

### PDMS Replica Molding

PDMS was made using a 10:1 ratio of PDMS base polymer and crosslinker by weight (Sylgard 184, Dow). After the PDMS base polymer and crosslinker were weighed, the mixture was continuously stirred by hand for 2 minutes. The PDMS was then poured into 3D printed molds and degassed in a vacuum desiccator (Bel-Art) for 15 minutes, the vacuum was then released and surfaced air bubbles were popped using a bulb air pump blower and desiccated for another 15 minutes. 3D printed molds with degassed PDMS were cured in a heated gravity convection oven. Curing temperatures were adjusted according to epoxy type, which have inherently different properties and temperature resistance: PDMS casted in molds coated with XTC-3D epoxy were cured at 40 °C for 4–5 hours, while PDMS casted in molds coated with Janchun were cured at 65 °C for 4–5 hours. After heat curing, the molds were allowed to cool at room temperature for 10 minutes before the PDMS was carefully peeled from the molds. As a heat-free alternative, PDMS can be cured at room temperature over 48 hours.

### PDMS Replica Characterization from 3D Printed Molds with Varied Geometries

3D printed molds with varied channel dimensions and geometric shapes were designed to evaluate the fidelity of PDMS replicas across a range of feature sizes. To assess the dimensional accuracy of PDMS replicas produced from 3D printed molds with and without epoxy coatings, we compared the replicated geometries against their original CAD designs. PDMS fluidic channels (nominal dimensions: 20 mm length, 1 mm width, 1 mm height) were fabricated from SLA and FDM 3D printed molds under different epoxy coating conditions. SLA printed molds were also fabricated for decreasing channel sizes ranging from 1 mm to 100 µm with cross-sectional widths and heights at a 1:1 aspect ratio, and with 20 mm channel length. All PDMS replica channels generated from FDM and SLA printed molds were characterized as follows: Channel lengths were measured using digital calipers with 10 µm resolution to ensure precision, while channel height and width were determined from brightfield images of cross-sectional slices acquired via an inverted microscope (Nikon Eclipse Ti2, 4X Objective). Cross-sections were analyzed in ImageJ, with measurements taken from the top, middle, and bottom regions of the molds to determine average dimensions.

A stacked rectangular structure, referred to as the “birthday cake” design, was fabricated using SLA printing. The structure consisted of three progressively smaller layers with widths of 3 mm (bottom layer), 2 mm (middle layer), and 1 mm (top layer), each with a uniform height of 1 mm. The birthday cake mold was coated with a Janchun epoxy 0.5 acetone mixture (1A:1B:0.5 acetone). As described previously, brightfield images of PDMS replica cross-sections were acquired, and feature dimensions were quantified using ImageJ. Measurements were obtained from the top, middle, and bottom regions of each layer to calculate the average width and height across three independent “birthday cake” structures. Measured values were compared to the original CAD design.

3D printed molds containing arrays of rectangular prisms and cylindrical post geometries were fabricated using SLA printing, with side lengths and diameters ranging from 1 mm to 300 µm, decreasing incrementally by 100 µm. Post heights were set at twice their corresponding side length or diameter. PDMS replicas were cast from 3D printed post molds coated with a Janchun epoxy 0.5 acetone mixture, producing rectangular and cylindrical cavities. The PDMS replicas were then placed in conformal contact with a glass coverslip, and brightfield images of the cavity openings were acquired. Average side lengths of square cavity faces and diameters of circular cavity faces were measured using ImageJ.

### Fluidic Device Fabrication

Cured PDMS was removed from 3D printed channel molds, cut into individual fluidic devices using a razor blade, and biopsy punched at channel edges to generate inlet and outlet holes. A 1.5 mm biopsy punch was used for tubing connections while a 6 mm biopsy punch created wells for gravity-driven flow. PDMS replicas were then sterilized in an ultrasonic cleaner with 70% ethanol (EtOH) for 15 minutes. Following sonication, parts were dipped into a beaker with sterile 100% EtOH to remove residual water droplets and allowed to dry in a biosafety cabinet. After drying, PDMS devices and glass coverslips were plasma treated by exposure to air plasma for 20 seconds using a plasma cleaner (PE-25, Plasma Etch). PDMS devices were then brought into contact with glass coverslips, placed on a hot plate at 65 °C, and left overnight to allow for complete sealing. To ensure maximum bonding strength while maintaining consistent conditions across fluidic devices generated from distinct molds, a glass plate holding a heavy beaker filled with water was placed on top of the PDMS-glass assemblies to apply uniform pressure during bonding. Bonded devices were plasma treated immediately before fluid introduction to render the channels hydrophilic. To assess leakage and confirm that channels were unobstructed, an aqueous solution dyed with red food coloring was introduced into one side of the devices, allowing gravity-driven flow from a defined height. Devices were inspected for immediate leakage and monitored over time for delayed fluid escape at the PDMS-glass interface or connection points. Fluid height was adjusted to apply pressure and identify potential sealing defects. Devices were observed under a light microscope at 10X magnification to detect leakage that may not have been visible to the naked eye.

### Visualization of Fluid Filled Interiors of Fluidic Channels by Fluorescent Dextran

PDMS channels with widths of 1000 µm, 800 µm, 600 µm, 400 µm, 200 µm, and 100 µm and a uniform 1:1 height to width ratio and total length of 20 mm were generated from SLA printed molds coated with Janchun epoxy 0.5 acetone mixture. Glass-bonded PDMS channels were prepared and rendered hydrophilic as previously described. Next, a 200 µL solution of Dextran conjugated to a fluorescent dye (Rhodamine B, 10,000 MW, Neutral; Thermo Fisher Scientific) was introduced into the channels. Confocal images were acquired with a 10X objective and a Z-step of 20 µm to visualize the fluorescent Dextran-Rhodamine dye (see **Microscopy methods**).

### Cell Culture

4T1 murine mammary carcinoma cells were cultured in RPMI-1640 medium with L-Glutamine (Corning) supplemented with a final concentration of 10% heat-inactivated fetal bovine serum (Gibco) and 1% gentamicin (Sigma). Here, we used E2 and M 4T1 clonal cell lines stably expressing near-infrared fluorescent protein (pNLS-iRFP670) and yellow fluorescent protein (YFP) in the nucleus, respectively, which we previously generated to enable live cell fluorescence imaging.^24^ pNLS-iRFP670 was a gift from Vladislav Verkhusha (Addgene plasmid # 45466; http://n2t.net/addgene:45466; RRID:Addgene_45466). The 4T1 E2 clonal cell line expressing pNLS-iRFP670 was routinely cultured in media containing 250 μg/mL G418 (Sigma) to apply selective pressure for the maintenance of pNLS-iRFP670 plasmid expression. 4T1 cells were routinely passaged using 0.05% trypsin/0.53mM EDTA in HBSS w/o Ca, Mg, or NaHCO_3_ (Corning). MCF10A human mammary epithelial cells were cultured in standard growth media established by the Brugge lab,^28^ consisting of DMEM-F12 with HEPES (Gibco) supplemented with final concentrations of 5% Horse Serum (Gibco), 20 ng/mL human epidermal growth factor (Gibco Human EGF, Animal-Free Recombinant Protein, PeproTech®), 0.5 µg/mL hydrocortisone (Sigma), 100 ng/mL cholera toxin (Sigma), 10 µg/mL insulin from bovine pancreas (Sigma), and 1% penicillin-streptomycin (Gibco). The MCF10A cell line used here is stably transfected with an inducible Snail expression construct fused to an ER response element, as well as green fluorescent protein (GFP) in the cytoplasm and red fluorescent protein in the nucleus (RFP-H2B), gifted by D.A. Haber.^29^ MCF10A cells were dissociated from the tissue culture flasks as a single-cell suspension using Accumax for routine passaging. All cell lines were cultured in an incubator at 37 °C and humidified atmosphere containing 5% CO_2_.

### Cell Viability Assay

Flat bottom molds with raised edges were 3D printed and coated with either XTC-3D or Janchun epoxy to generate a flat slab of PDMS to serve as a cell culture surface. Control PDMS slabs were prepared by directly pouring and curing PDMS in a 10 cm polystyrene dish, which served as a biocompatible mold without an epoxy coating. Cured PDMS replicas were then removed from their respective molds, trimmed to remove the curved edges and obtain a ~ 7 cm × 7 cm slab, and subsequently sterilized in an ultrasonic cleaner bath with 70% ethanol (EtOH) for 15 minutes.

After sonication, parts were dipped into a beaker with sterile 100% EtOH to remove any residual water droplets and allowed to dry in a biosafety cabinet. Dry, sterilized PDMS replicas were then placed in a 10 cm polystyrene dish and coated with Type 1 collagen derived from rat tail (Corning) at 5 µm/cm^2^. To enhance collagen adhesion to the hydrophobic PDMS surfaces, the PDMS replicas were exposed to plasma for 20 seconds in a plasma cleaner. After plasma treatment, a dilute collagen solution prepared in 0.02N acetic acid was gently spread over the plasma treated PDMS surface and incubated at room temperature for 1 hour in a biosafety cabinet, avoiding any disturbances that could break the surface tension.

After the incubation period, the excess collagen was aspirated, and the PDMS slabs were washed three times with 1x PBS prior to seeding 4T1 and MCF10A cells on the collagen coated surfaces. Cells were seeded at a concentration of 800K cells per PDMS slab by suspending cells in 5 mL of media, which was carefully deposited onto the PDMS molds, ensuring an even distribution while maintaining the integrity of the surface tension. The culture dishes were subsequently placed in an incubator for 1 hour to allow the cells to adhere to the PDMS surface, followed by the addition of media to the dishes resulting in fully submerged PDMS cultures.

The cell monolayers were examined after 3 days of incubation, when cell confluence reached ~ 80%, at which point brightfield and epifluorescence images of the live cell cultures were acquired using an inverted microscope and a 20X objective (see **Microscopy methods**). Following imaging, cell viability was assessed by trypsinizing cells and performing a trypan blue exclusion assay with a final concentration of 0.2% trypan blue (Cytiva HyClone). The stained cell suspension was then loaded into a hemocytometer, and both viable (unstained) and nonviable (blue stained) cells were manually counted under an inverted tissue culture microscope (Nikon Ts2 inverted microscope) with a 10x phase contrast objective (CFI ACHRO ADL 10x PH1 NA 0.25 WD 5.2MM). Cell viability was calculated as the percentage of live cells in the total cell count [(live cells/live + dead cells) × 100].

### Generation of 3D Tissue Aggregates in Fluidic Channels

PDMS fluidic channels (nominal dimensions: 20 mm length, 1 mm width, 1 mm height) were generated from SLA printed molds, sterilized, bonded to glass coverslips, and rendered hydrophilic via plasma treatment, as described. Next, 3D cell aggregates were engineered in a type 1 collagen matrix housed within these fluidic channels, as previously described.^30^ Briefly, a sterile solution of 1 mg/mL Poly-D-lysine (PDL) (A-003-E, Sigma-Aldrich) was injected into the fluidic PDMS channels following plasma treatment and left in a biosafety cabinet for 1 hour to coat the interior of the channels. The PDL solution was aspirated from the channels, which were then washed three times with sterile filtered type 1 ultra-pure water. Acupuncture needles with 120 µm diameter were coated with a 1% sterile solution of bovine serum albumin for 1 hour at room temperature and subsequently washed three times with ultra-pure water. Coated acupuncture needles were then aligned in the center of PDL coated fluidic channels and the devices were cooled to 4 °C before proceeding to the following step.

Next, a 4 mg/mL type 1 collagen gel was molded around the acupuncture needle within the fluidic channel to generate a hollow collagen cavity. PDMS was precured by heating at 65 °C for 10 minutes to be used for sealing the devices. First, a small amount of precured PDMS was place at the open end of the channel to plug the channel. Next, chips were placed over the open window region of 3D printed imaging plates and precured PDMS was used to seal the coverslip to the plate bottom. The devices mounted on 3D printed imaging plates were then placed in the incubator overnight at 37 °C to allow PDMS to fully cure. Subsequently, 4T1 M clonal cells with nuclear YFP expression, and MCF10A cells with cytoplasmic GFP expression and RFP-H2B nuclear expression, were seeded into distinct cavities to form solid 3D cell aggregates, mimicking breast tumor and breast tissue aggregates, respectively. Confocal fluorescence images of 3D cell aggregates were acquired three days after cell seeding on an inverted laser-scanning confocal microscope using a resonant scanner and 20X objective (**Microscopy Methods**).

### Organ-on-a-Chip Platform: Inventory of Parts and Dimensions

A custom 3D printed imaging platform was designed to match the dimensions of a standard 96-well plate to ensure compatibility with existing laboratory equipment. The device dimensions are 126.56 mm × 85 mm. The design was created using SOLIDWORKS and printed using a Formlabs Form3+ with Clear V4 resin. The platform wall was extruded to a total height of 8.45 mm, designed to accommodate a universal well plate lid. To allow for direct imaging of fluidic devices using an inverted microscope, the platform incorporated four rectangular openings (60.5 mm × 24.5 mm) designed to securely hold 60 mm × 24 mm glass coverslips. Each opening included a 1 mm overhang along its perimeter to prevent the coverslips from dislodging during handling. To maintain a hydrated microenvironment suitable for cell culture imaging, two reservoirs were incorporated at both the top and bottom of the platform. These reservoirs were designed to hold either 1X phosphate buffered saline or water, helping to prevent evaporation and maintain stable culture conditions. The platform was tested on a Nikon Eclipse Ti2 microscope equipped with a stage top incubator to ensure compatibility for live-cell imaging and the use of air and immersion objectives.

Next, A 3D printed mold for PDMS chamber arrays was designed using SOLIDWORKS and manufactured using the Form 3+ with Clear V4 resin. The mold dimensions were 125 mm × 75 mm, with a thickness of 10 mm. For ease of demolding, the mold’s edges were filleted with a 4 mm radius, and the top face was shelled to a thickness of 2 mm. The mold contains a 4 × 5 array of channels, with each individual channel measuring 1 mm in height and width, and 20 mm in length. The spacing between channels was designed to facilitate PDMS cutting and handling: within each column of 5 channels, channels were spaced 9.5 mm apart, while the distance between adjacent columns was 10 mm to allow for precise cutting.

Finally, a 3D printed mold comprising an array of posts was designed using SOLIDWORKS to fabricate PDMS wells. The mold dimensions were 80 mm × 70 mm, with a thickness of 10 mm. For ease of demolding, the mold’s corners were filleted with a 4 mm radius, and the top face was shelled to a thickness of 2 mm. An array of 4 × 5 cylindrical posts was incorporated into the mold, each with a diameter of 8 mm and an extrusion height of 4 mm. The distance between each post is 2.5 mm for a strip of 5 posts and the distance between 4 posts is 5 mm for ease of cutting. This design was intended to produce uniform PDMS wells consistent with the geometry and dimensions of the chamber arrays for the OoCs. The fabrication and testing of all 3D printed parts for the OoC imaging platform were performed in 3 technical replicates each (n = 3) to ensure reproducibility and validate the platform’s structural integrity and functionality.

### Microscopy

All acquired images were of live cell cultures. Brightfield and epifluorescence images were acquired on the Nikon Eclipse Ti2 microscope with a D-LEDI Fluorescence LED Illumination System equipped with an ORCA-FLASH 4.0 LTS+ SCMOS camera. The filter sets used to image red, green, and near-infrared fluorescent proteins included C-FL GFP HC HISN Zero Shift, C-FL DS RED HC HISN Zero Shift, and C-FL CY5 HC HISN Zero Shift, respectively. Imaging was performed using either a CFI Achromat Flat Field 4X objective, CFI60 Plan Apochromat Lamda D 10X objective, or CFI60 Plan Apochromat Lamda D 20X objective, as indicated. Confocal fluorescence images were acquired on the Nikon AX R laser scanning confocal microscope system equipped with a LUA-S4 laser unit, which provided excitation at 488 nm and 561 nm wavelengths. Emission was captured using 521/42 and 600/45+650LP filters for yellow and green or red fluorescent proteins, respectively. Confocal fluorescence images were acquired using a resonant scanner and CFI Apochromat Long Working Distance Lambda S 20X Water Immersion objective. All acquired images were of live cell cultures.

### Statistical Analysis

Statistical significance was determined using a two-tailed t-test for comparison of two conditions for the cell viability assays. Statistical significance was determined using a one-way ANOVA test for comparison of three or more groups for PDMS replica characterization, which was followed by a post-ANOVA Tukey test to assess pairwise differences between groups. Bar plots show the values for average measurements and error bars represent the standard deviation of the corresponding data. To aid in the visualization of significance between many experimental conditions, we generated pairwise significance heatmaps, where colors and asterisks indicate the level of statistical significance: blue = not significant; gray/* = p-value < 0.05; pink/** = p-value < 0.01; red/*** = p-value < 0.001 (**Fig. S1, Fig. S3**). Three independent experiments were conducted for all cell studies. To account for potential variability in 3D printing and epoxy coating conditions, five distinct 3D printed structures were measured to characterize PDMS replicas derived from 3D printed molds.

## Results

### Optimizing Epoxy Coatings for High-Fidelity PDMS Replication from SLA Printed Molds

To overcome the limitations associated with surface roughness and poor feature fidelity of PDMS replicas generated from 3D printed molds, we established a robust 3D printing post-processing method that enables reliable PDMS replica fabrication and enhances compatibility with downstream PDMS-based fluidic applications. The novel method produces reusable molds and is compatible with both SLA and FDM printing (**Figure 1**).

**Figure 1.**
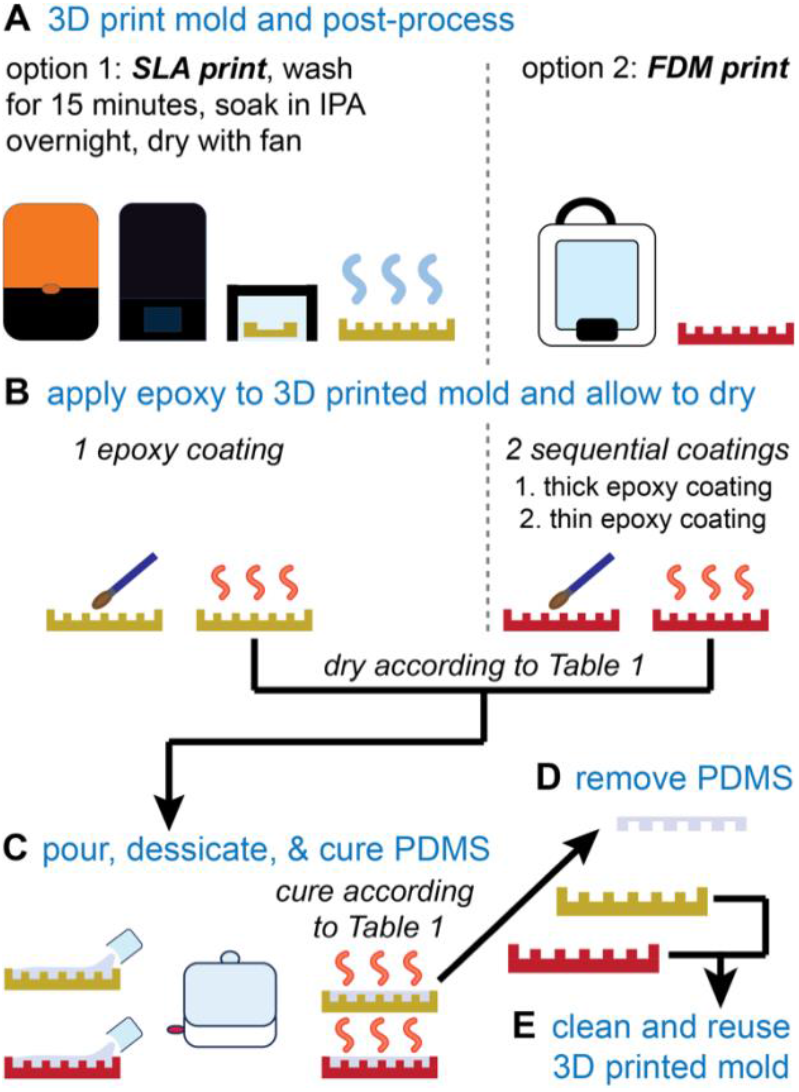
3D print post-processing workflows for 3D printed mold fabrication. **(A)** 3D printing, washing, and drying steps for SLA and FDM printed molds. **(B)** Application and drying of epoxy coatings on SLA and FDM printed molds based on Table 1 specifications. **(C)** Pouring, desiccation, and curing of PDMS based on Table 1 specifications. **(D)** Removal of cured PDMS replicas from SLA and FDM printed molds. **(E)** Cleaning and reuse of 3D printed molds.

We hypothesized that applying an epoxy coating to 3D printed molds would improve surface quality, facilitating repeated and high fidelity PDMS replica molding. To test this, we first designed and fabricated 3D printed molds with diverse millifluidic channel sizes and geometric shapes using a Form 3+ SLA printer. We chose to evaluate the XTC-3D system for coating 3D printed molds because it is specifically marketed as a protective coating to smooth and finish 3D printed parts, and it is compatible with a wide range of 3D printing materials used for SLA and FDM printing. Following the fabrication and standard post-processing of the 3D printed parts, we applied a thin layer of XTC-3D epoxy, carefully coating the surface that would serve as the template region for replica molding. To do this, we prepared an epoxy mixture of XTC-3D at a 2:1 ratio of part A (resin) to part B (hardener) and brushed this on to the 3D print surface using a small flat paintbrush, following the manufacturer’s recommendations. After the epoxy coating had completely dried, PDMS precursor was poured onto the 3D printed molds and allowed to cure. PDMS replicas were then carefully removed, sterilized, and bonded to clean glass coverslips. The resulting PDMS devices exhibited strong bonding, evidenced by resistance to mechanical removal, the ability to withstand fluid injection into the hollow regions of the devices formed by the raised features of the 3D printed molds without leakage, and no signs of delamination (**Figure 2**). To demonstrate the versatility of the fabrication approach, we created devices with chambers featuring various geometries, including right angle shapes, bifurcated structures, serpentine channels, and custom shapes like a pentagon and a star (**Figure 2A**). To showcase the fluidic capabilities of fabricated devices, we embedded continuous channels that connect to tubing for fluidic control. We generated devices with intricate, spatially varying channels, such as the ‘ILLINI’ pattern, and decreasing channel widths to demonstrate flexibility in accommodating both complex designs and fluidic flow (**Figure 2B**). These examples served as a proof of principle, illustrating the ability of epoxy-coated SLA 3D printed parts to serve as molds for fabricating customizable PDMS devices suitable for millifluidic applications.

**Figure 2.**
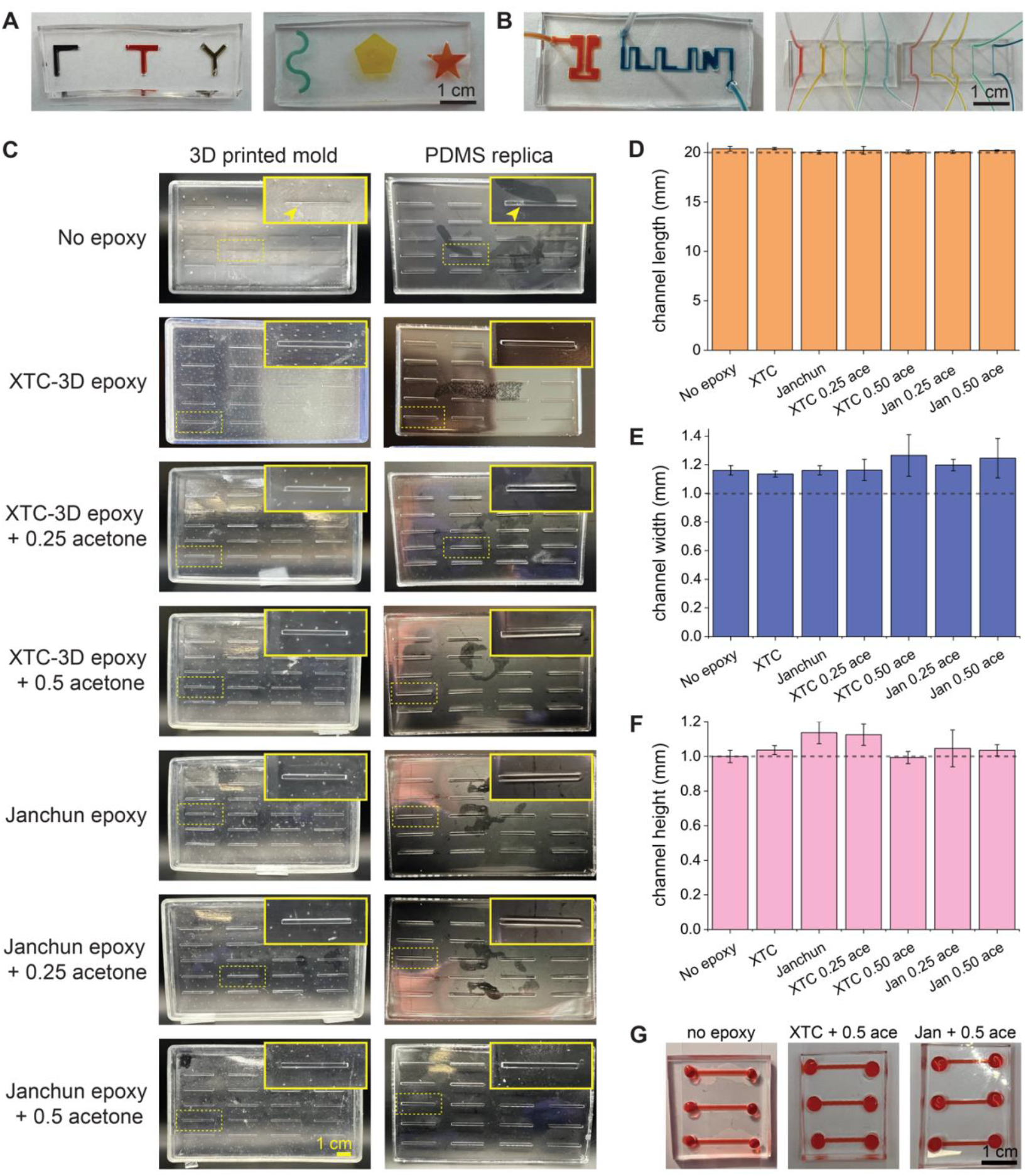
Diverse geometries and characterization of PDMS-based devices. **(A-B)** PDMS-based devices fabricated from 3D printed molds coated with XTC-3D epoxy. **(A)** Millifluidic T-junctions and Y-junctions, as well as serpentine, pentagon, and star designs were cast to show the geometric complexity achievable with this method. **(B)** Millifluidic “ILLINI” device with embedded tubing to demonstrate fluidic connectivity. Channels with decreasing cross-sectional dimensions ranging from 1 mm to 100 µm were fabricated to showcase size scalability. **(C)** PDMS channels molded from SLA printed molds. The molds were coated with either XTC-3D or Janchun epoxy, with or without the addition of acetone, as indicated. The top row shows an uncoated mold as a baseline control. Yellow insets highlight surface smoothness and potential defects transferred from the mold to the PDMS replica, showing how epoxy coatings impact mold quality and replication performance. **(D-F)** Quantification of channel **(D**) length, **(E)** width, and **(F)** height of PDMS replicas fabricated from molds with varied epoxy coating conditions; Bar graphs show mean and standard deviation, and the dashed line indicates nominal CAD dimensions. **(G)** PDMS devices bonded to glass coverslips were filled with red food dye to evaluate optical clarity and sealing performance. devices cast from epoxy-coated molds exhibited strong bonding and no leakage, whereas devices from uncoated molds showed poor transparency, failed bonding, and extensive leakage. Abbreviated terms represent XTC-3D (XTC), Janchun (Jan), and acetone (ace). Scale bars = 1 cm.

Next, we aimed to conduct a more comprehensive evaluation of epoxy coating 3D printed molds for PDMS replica molding. Here, we generated 3D printed molds containing 20 channels with nominal channel dimensions of 20 mm in length, 1 mm in width, and 1 mm in height based on CAD designs. Channels were arranged across 4 columns, each with 5 channels and equidistant spacing across a row (**Figure 2C**). This configuration was chosen for downstream applications to yield 4 PDMS devices with 5 fluidic channels each. SLA printing parameters were set to 100 µm resolution generating 3D printed molds with moderate striations. We hypothesized that thinned epoxy mixtures may provide an optimal surface coating to fill these minor grooves, while minimizing the thickness of the epoxy coating that could increase feature size. Accordingly, 3D printed molds were coated with either no epoxy, standard epoxy, or thinned epoxy produced by the addition of acetone to epoxy mixtures. We first painted epoxy on the 3D printed mold surface, then applied a controlled stream of air to precisely spread the epoxy across the mold features, ensuring a smooth, finish. A standard mixture of XTC-3D epoxy was prepared per the manufacturer’s instructions, combining 2 parts A and 1 parts B. To thin the epoxy to different extents, we added acetone at different ratios, including a thin 0.25 acetone mixture (2 parts A, 1 parts B, and 0.25 parts acetone), and thinner 0.50 acetone mixture (2 parts A, 1 parts B, and 0.50 parts acetone). Moreover, we aimed to evaluate the suitability of alternative epoxy systems for coating 3D printed molds. Since XTC-3D is a two-part system with a 2:1 resin to hardener ratio, we chose to evaluate a two-part system with a 1:1 ratio using JANCHUN Crystal Clear Epoxy Resin, which we refer to as ‘Janchun’ epoxy. Similar to XTC-3D coated 3D printed parts, Janchun epoxy was applied as a standard epoxy mixture (1 part A, 1 part B), as well as with a 0.25 acetone mixture and 0.50 acetone mixture, representing the addition of 0.25 parts acetone and 0.50 parts acetone, respectively. Epoxy coating dry times were optimized for room temperature and elevated temperatures (**Table 1**). Elevated temperatures significantly reduced drying times from 12 hours to 1–2.5 hours. Molds coated with Janchun epoxy cured at 65 °C within 1.5 hours, while molds coated with XTC-3D epoxy cured at 40 °C within 2.5 hours. We optimized the drying times at selected temperatures based on the manufacturer’s recommended curing range, focusing on the midpoint temperature to minimize risks associated with higher temperatures, such as melting or deformation. Complete epoxy curing was verified following standard physical and chemical evaluation (**Methods**). Optimal drying times were determined based on epoxy type and acetone content, with higher acetone concentrations requiring extended drying durations (**Table 1**). Figure 2C highlights the differences in surface quality and transparency of PDMS replicas cast from molds coated with no epoxy, XTC-3D epoxy, and Janchun epoxy mixtures. Uncoated molds with no epoxy yielded PDMS replicas with an opaque appearance, poor transparency, and noticeable irregularities, with sites of PDMS tearing and incomplete removal from molds. In contrast, epoxy-coated 3D printed molds yielded PDMS replicas with significant improvements in quality and clarity across all epoxy conditions compared to uncoated molds. We found that PDMS replicas generated from 3D printed molds coated with epoxy mixtures containing acetone appeared smoother and clearer compared to molds coated with epoxy without acetone. For both XTC-3D and Janchun epoxy, the best overall results were obtained for 3D printed molds coated with 0.5 acetone mixtures, yielding PDMS replicas with minimal surface imperfections and notably improved transparency (**Figure 2C**). This improvement can be attributed to the acetone’s ability to reduce the viscosity of the epoxy mixture, enabling it to penetrate and smooth surface irregularities during curing. To quantify the channel fidelity for PDMS replicas obtained from different 3D printed molds, we measured channel lengths, widths, and heights across epoxy treatment conditions compared to a control, uncoated 3D printed mold (**Figure 2D-E**). The channel length remained consistent across all treatment groups, with no significant deviations from the nominal 20 mm length (**Figure 2D, Figure S1A**). Due to the inability to directly measure channel width and height, measurements were made based on brightfield images acquired for cross-sectional slices of the PDMS channels. No significant differences in channel width were observed; However, measured channel widths were ~200 µm higher than the nominal 1 mm CAD design across all conditions, with or without an epoxy coating (**Figure 2E, Figure S1B**). The discrepancy between nominal versus actual measured width may be explained by the challenge in placing channel cross-sections on their side for optical imaging and subsequent measurement of cross-sectional dimensions. In contrast, channel heights more closely matched the nominal 1mm distance across all conditions (**Figure 2F**). Statistically significant increases in channel height were observed for the XTC-3D 0.25 acetone mixture and Janchun epoxy compared to the control uncoated mold, as well as compared to the XTC-3D 0.5 acetone mixture (**Figure S1C**). This result is expected, as the deposited epoxy layer could naturally thicken the features of the 3D printed mold. Notably, epoxy mixtures with 0.50 acetone produced mean channel heights that most closely matched the nominal 1 mm CAD design for both the XTC-3D and Janchun epoxy systems (**Figure 2F**). The enhanced height fidelity observed for epoxy-acetone 0.50 mixtures is likely due to the optimal balance of viscosity and surface penetration, allowing for uniform epoxy application and minimal vertical distortions. These results demonstrate that incorporating acetone into epoxy mixtures, particularly at a 0.50 acetone ratio, enhances the surface quality, optical clarity, and dimensional fidelity of the resulting PDMS channels compared to uncoated molds and epoxy-coated molds with little or no acetone.

Based on these findings, we primarily focused on epoxy-acetone 0.50 mixtures for subsequent characterization of epoxy-coated molds and PDMS replicas. To compare the performance of fluidic channels generated from molds with and without an epoxy coating, we punched inlet and outlet holes at the ends of representative PDMS channels and plasma bonded the devices to clean glass coverslips. Next, we added a dilute solution of red food coloring dye in water at one end of the channels and examined fluid flow through and retention in the channels (**Figure 2F**). We observed that the red solution was able to completely flow through channels from the inlet to outlet for all devices; However, the solution exhibited immediate leakage across the entire PDMS-glass interface for channels generated from the uncoated 3D printed mold, with signs of delamination at channel edges. Moreover, these devices could be readily removed from their underlying glass coverslips, indicating plasma bonding was unsuccessful. In contrast, no fluid leakage was observed for channels generated from 3D printed molds coated with 0.5 acetone mixtures of either XTC-3D or Janchun as the epoxy base (**Figure 2F**). Unlike PDMS replicas from uncoated molds, PDMS replicas from coated molds did not exhibit delamination in any regions of the devices and could not be physically removed from their underlying glass coverslips, indicating successful and irreversible plasma bonding. Overall, these findings underscore the potential of epoxy-coated 3D printed molds as a versatile and effective approach for producing high quality PDMS replicas for fluidic applications.

### A Robust Approach for Transparent PDMS Replicas via Acetone-Thinned Epoxy-coated 3D Printed Molds

Given our findings that 3D printed molds coated with epoxy-acetone 0.50 mixtures yielded optimal PDMS replicas, the next phase of our work focused exclusively on this optimized formulation to further evaluate its capability for fabricating 3D structures and their effective applications. Given the considerable visible improvement in PDMS transparency with epoxy-coated molds, we expected that these devices would be highly suited to optical microscopy applications (**Figure 3**). To demonstrate this, we acquired brightfield images of the channel regions for PDMS devices placed on glass coverslips, taking an image through an anchored device (top view) and image of a channel cross-section that was cut and placed on its side. Images of PDMS devices obtained from uncoated 3D printed molds revealed a rough and irregular surface at the top of the channel, obstructing optical imaging through the hollow region of the channel. Images of channel cross-sections depict inconsistencies in channel walls, exhibiting wrinkle-like features at channel edges (**Figure 3A**). These undesirable features can be attributed to the noticeable stair-stepping artifacts from the SLA printing process and PDMS defects from tearing at the mold interface (**Figure 2C**). In contrast, images of PDMS devices from XTC-3D epoxy-coated molds revealed a smooth surface at the top of the channel, which was associated with superior optical imaging throughout the PDMS (**Figure 3A**). Cross-sectional views further highlight this improvement, with epoxy-coated molds producing more uniform channel profiles that closely matched the CAD designs. PDMS replicas from Janchun epoxy-coated molds achieved the best results, demonstrating exceptional optical clarity and complete absence of visible artifacts at channel walls (**Figure 3A**). Accordingly, to examine the breadth of our fabrication approach, we designed and generated 3D printed molds coated with Janchun epoxy-acetone 0.50 mixtures for molds with a sweep of geometries and size-scales. As previously discussed, milliscale features are challenging to obtain with standard lithography approaches, especially for complex multilayered geometries. As such, we designed and fabricated a three-tier “birthday cake”-like structure fluidic channel with distinct dimensions for the base, middle, and top tiers. The CAD design specified a 1 mm height for each layer µm, and tier widths of 3 mm for the base layer, 2 mm for the middle layer, and 1 mm for the top layer. The resulting PDMS replicas yielded from these “birthday cake” molds successfully preserved the overall geometry of the tiered structure, with clear and distinct transitions between the base, middle, and top tiers (**Figure 3B**). Quantitative measurements of individual tier dimensions indicated that the replica heights closely matched the CAD design, while slight increases were observed for the tier widths (**Figure 3C**). These deviations are likely due to epoxy buildup during coating of the SLA printed mold during curing.

**Figure 3.**
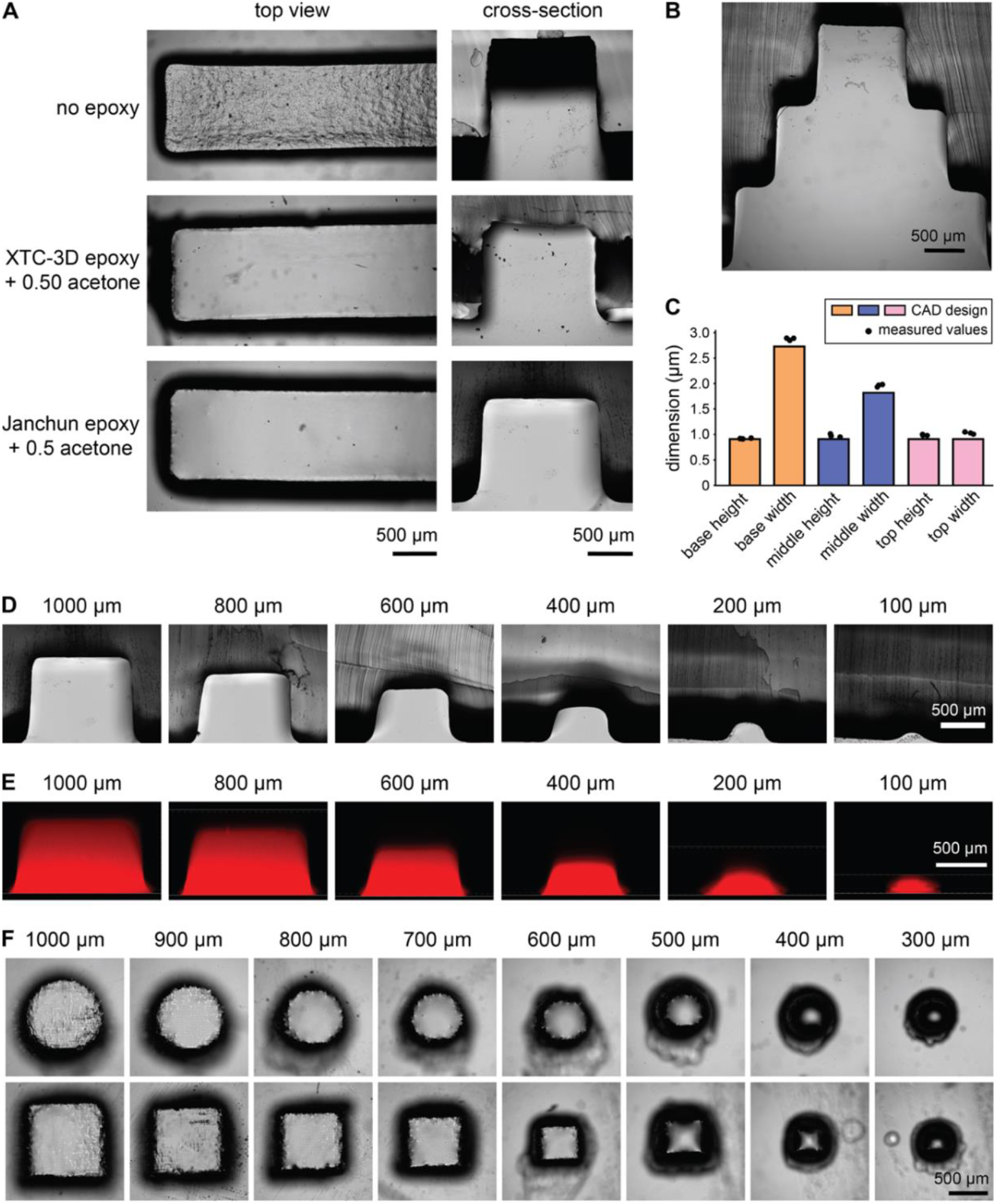
Fidelity of PDMS replication using molds coated with acetone-thinned epoxy. **(A)** Brightfield images of top and cross-sectional views of PDMS channels fabricated using molds under three conditions: no epoxy coating, XTC-3D epoxy with 0.50 acetone, and Janchun epoxy with 0.50 acetone. Uncoated mold yield PDMS replicas with a rough surfa ce, whereas epoxy-coated molds yield smooth, well-defined channels. **(B)** Brightfield image of a PDMS replica fabricated from a mold with a “Birthday cake” structure, demonstrating the ability to replicate multi-tiered geometries with layer widths from 1 mm to 3 mm. **(C)** Bar graph illustrating how the measured dimensions of the “birthday cake” compares to its original CAD design. **(D-E)** PDMS channels fabricated from molds with decreasing channels sizes, ranging from 1000 µm to 100 µm. Brightfield images of channel cross-sections show channel walls **(D)** and fluorescent confocal microscopy images of PDMS channels bonded to glass coverslips and filled with a Dextran-rhodamine dye to visualize the channel fluid-filled interior **(E)**; Images show that channel cross-sectional shape begins to shift from square to more rounded for width and height of <400 µm, while channels remain open and retain fluidic capacity down to 100 µm dimensions. **(F)** Circular and square cavity replicas with diameter and side lengths from 1000 µm to 300 µm, revealing progressive loss of structural definition at smaller sizes, with square features maintaining slightly better shape fidelity than circular ones at 400 µm. All corresponding molds were coated with Janchun epoxy-acetone 0.50 mixtures.

To further evaluate the fidelity and functionality of our fabrication process, molds with decreasing channel sizes ranging from 1 mm to 100 µm were generated with cross-sectional widths and heights at a 1:1 aspect ratio, and with 20 mm channel length. A thin cross-sectional slice was cut from a subset of PDMS replicas, and brightfield images were acquired to confirm that channels were open at the center of the devices (**Figure 3D**). A separate subset of devices was bonded to glass coverslips to examine fluid flow through the channels. To do this, a solution of dextran conjugated to fluorescent Rhodamine dye was first introduced into a single well of the bonded devices, leaving the opposing well empty. The dextran solution visibly filled the entire channels, reaching the opposing well via gravity-driven flow. Confocal imaging of the fluorescent dextran was conducted to examine the fluid distribution within the channels (**Figure 3E**). Confocal images were acquired as Z-stacks with a step size of 20 µm, revealing a fluorescent channel profile with square edges in channels with cross-sectional dimensions ranging from 400 µm to 1 mm, indicating excellent replication fidelity and structural integrity. Notable distortions were observed in channels with cross-sectional dimensions of ≤200 µm, exhibiting visible rounding of the channel walls and a shift in the fluorescent profile toward half-cylindrical geometries (**Figure 3D-E**). These results highlight the advantage of epoxy-coated 3D printed molds for generating functional PDMS channels that support fluid flow across a wide range of cross-sectional dimensions. This approach ensures reliable channel formation and fluid transport, making it well-suited for applications requiring precise fluidic control.

To examine geometric versatility and resolution limits beyond fluidic channel designs, we fabricated epoxy-coated 3D printed molds with arrays of posts with decreasing feature sizes from 1 mm to 300 µm, representing the side lengths and diameters of square and circular faces, respectively. Post heights were set to either equal (1:1) or twice (2:1) their corresponding side length or diameter, yielding PDMS replicas with cylindrical and rectangular prism cavities of decreasing feature size. Characteristic circular and square post faces remained well-defined down to at least 500 µm (**Figure 3E**) for 2:1 geometries. Square-faced cavities closely followed the dimensions of their CAD designs, while circular-faced cavities exhibited greater shrinkage, especially for smaller feature sizes in the 1:1 aspect ratio (**Figure S2**). These findings highlight the impact of aspect ratio on feature fidelity in our approach, demonstrating that a 1:1 aspect ratio can effectively produce high fidelity fluidic channels down to at least 400 µm, while increased aspect ratios may be required for similarly side post features in a shape dependent manner.

Together, these results demonstrate the versatility of epoxy-coated 3D printed molds for fabricating high fidelity PDMS channels that support fluid flow across a broad range of cross-sectional dimensions, spanning an entire order of magnitude from 1 mm down to 100 µm. Additionally, this approach enables the fabrication of replica molded prism structures with tunable aspect ratios, optimizing feature fidelity for microstructured array applications and expanding its utility for microscale patterning and structured well designs. These capabilities highlight the adaptability of our method, particularly excelling for larger microfluidic and millifluidic applications.

### Optimizing Epoxy Coating on FDM Printed Molds for Fluidic Applications

To assess the compatibility of our coating method with a more cost-effective 3D printing approach, we investigated FDM for fabricating fluidic molds. Compared to SLA, FDM offers significantly lower material costs and a simplified workflow; However, its lower resolution and characteristic layer striations can result in surface imperfections and reduced feature fidelity of 3D printed parts. Here, we fabricated FDM printed molds using PLA filament, reproducing the same 20 channel CAD design as the SLA printed molds (**Figure 2, Figure 4**). PDMS replicas generated from untreated FDM printed molds suffered from significant damage and visible striations. Notably, small pieces of PDMS remained adherent to molds following replica removal, resulting in severe tearing at channel sites (**Figure 4a**).

**Figure 4.**
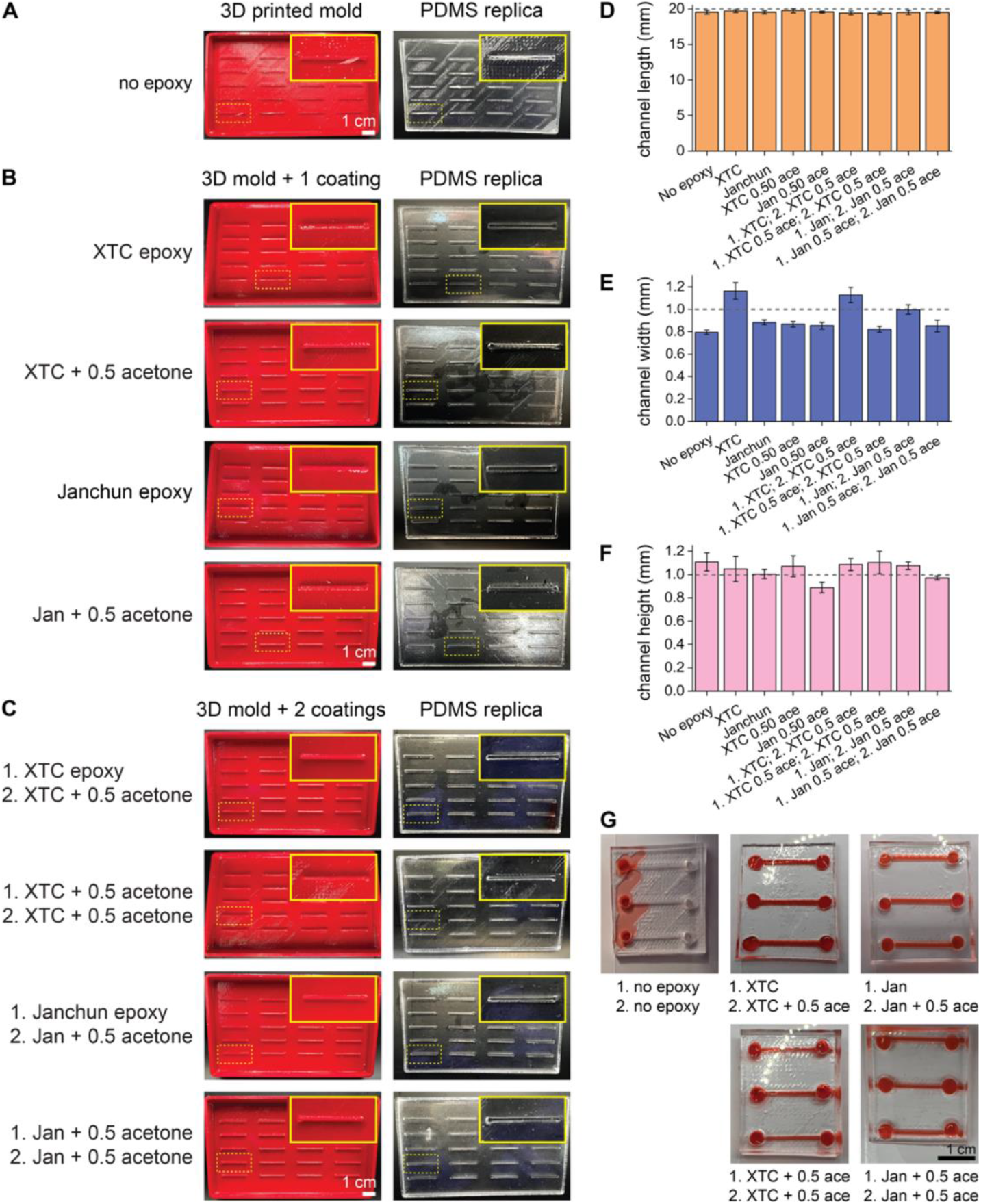
Two-step epoxy coating improves PDMS replication from FDM printed molds. **(A)** Comparison between an uncoated FDM printed mold (left) and its corresponding PDMS replica (right); The uncoated mold shows significant surface roughness and defects, which are transferred to the PDMS replica. **(B)** Evaluation of FDM printed molds with a single layer of epoxy coating **(B)** or double layer coating **(C)**, with XTC-3D epoxy, XTC-3D + 0.5 acetone, Janchun epoxy, or Janchun + 0.5 acetone. **(B)** A standard epoxy mold coating generates PDMS replicas with improved surface smoothness and more well-defined channels compared to acetone-thinned epoxy coatings. **(C)** Double epoxy-coated molds further improve channel definition of PDMS replicas, reducing roughness and enhancing the clarity of fluidic channels, with best results obtained with a first coating of standard epoxy followed by a second coat of a thinned-epoxy mixture. **(D-F)** Quantification of channel **(D**) length, **(E)** width, and **(F)** height of PDMS replicas fabricated from molds with varied epoxy coating conditions; Bar graphs show mean and standard deviation, and the dashed line indicates nominal CAD dimensions. **(G)** PDMS devices bonded to glass coverslips and filled with red food dye demonstrate bonding performance and fluid containment. Devices cast from uncoated molds exhibit poor transparency, failed bonding, and major leakage, whereas replicas from two-step epoxy-coated molds show strong bonding, optical clarity, and no leakage. Abbreviated terms represent XTC-3D (XTC), Janchun (Jan), and acetone (ace). Scale bars = 1 cm.

To address these challenges, we tested various epoxy coating strategies to enhance the quality and functionality of PDMS replicas generated from FDM printed molds. Based on our findings from SLA printed molds, we first assessed the suitability of either standard epoxy or epoxy-acetone 0.50 mixtures as mold coatings. We found that PDMS replicas could be cleanly removed from molds coated with standard epoxy mixtures for both XTC-3D and Janchun epoxy, yielding intact channels free of PDMS tears. In contrast, while molds coated with epoxy-acetone 0.5 mixtures produced PDMS replicas with higher fidelity than uncoated molds, these coatings poorly filled the grooves of FDM printed molds, leaving deep striations in both the coated molds and the resulting PDMS replicas (**Figure 4b**). Given the overall benefit of applying a single epoxy layer, we next investigated whether a second coating could further improve the PDMS replica molding process. Accordingly, we implemented a two-step epoxy coating process in which precoated molds (1. standard epoxy mixture or 1. epoxy-acetone 0.50 mixture) were coated with a second layer of epoxy (2. epoxy-acetone 0.50 mixture). We chose to apply the epoxy-acetone 0.50 mixture as the second coating to prevent excessive thickening of mold features, which would occur with two layers of standard epoxy. The second epoxy layer further smoothed the FDM printed mold surfaces, producing a high gloss finish for molds initially coated with the standard epoxy mixture and better groove filling for molds initially coated with the epoxy-acetone 0.50 mixture, which further improved the PDMS replicas (**Figure 4c**). Measurements of PDMS channel dimensions showed no significant differences in channel lengths (**Figure 4d, Figure S3a**), while cross-sectional widths and heights varied across epoxy coating conditions (**Figure 4e-f, Figure S3b-c**). Notably, most coating conditions produced channels with mean widths below the intended 1 mm CAD design, likely due to surface and mechanical forces during the replica molding process in molds with rougher surfaces (e.g., uncoated or thinly coated FDM molds), yielding suboptimal results. In contrast, relatively wider channels were observed in PDMS replicas from FDM printed molds coated with either a single layer of standard XTC-3D epoxy or a two-step coating where standard epoxy (XTC-3D or Janchun) was used as the first layer (**Figure 4e, Figure S3b**). Channel heights were similar across most conditions, with mean values near or slightly above the CAD dimension of 1 mm. The exception was molds coated with a single layer of Janchun epoxy-acetone 0.50 mixture, which yielded significantly shorter PDMS channels (**Figure 4f, Figure S3c**). This reduction in height may result from pooling of the thinned epoxy mixture at the mold base rather than evenly coating the entire mold and channel features. Next, we generated PDMS-glass devices by plasma bonding to examine how PDMS replicas from two-step epoxy-coated molds perform compared to uncoated FDM printed molds. Unsurprisingly, PDMS replicas from uncoated molds were found to be unsuitable for fluidic applications, exhibiting poor PDMS-glass bonding and immediate fluid leakage at the interface (**Fig. 4G**). In stark contrast to the defective PDMS replicas from uncoated molds, replicas generated from two-step coated molds exhibited irreversible PDMS-glass bonding and maintained fluid within the channel interiors **(Fig. 4G)**. As such the two-step epoxy coating process dramatically improved the surface quality of PDMS replicas compared to uncoated FDM printed molds, making them suitable for fluidic applications. Overall, the optimal FDM printed molds were obtained through two-step epoxy coating with 1. a standard epoxy mixture and 2. an epoxy 0.5 acetone mixture, yielding smooth surfaces and PDMS replicas with high fidelity channels. These findings demonstrate that low cost FDM printed molds, when properly coated, can serve as viable alternatives to SLA printed molds for fluidic device fabrication.

### Biocompatibility of PDMS Replicas from Epoxy-coated 3D printed Molds

Having established a robust epoxy coating method for FDM and SLA printed molds, we next evaluated the biocompatibility of PDMS replicas for biological applications. To do this, We SLA printed molds with a flat bottom and raised edges, applying a single coat of 0.5 acetone mixture (XTC-3D or Janchun), previously optimized for SLA prints (**Figure 2–3**). These molds served to generate flat slabs of PDMS for use as 2D cell culture surfaces. As a control, we molded PDMS in a Petri dish, an inherently biocompatible substrate with a flat surface. Cured PDMS slabs were removed from their respective molds, sterilized in 70% ethanol, and coated with type I collagen prior to seeding cells. Next, we cultured normal (MCF10A) and malignant (4T1) mammary epithelial cells expressing endogenous fluorescent proteins on the collagen coated PDMS slabs, examining readouts of biocompatibility at a 72 hour endpoint (**Figure 5**). The cell lines used here express endogenous fluorescent proteins, with GFP in cytoplasm and RFP in the nucleus of MCF10A cells, and near infrared fluorescent protein in the nucleus of 4T1 cells, enabling fluorescence imaging of cell cultures. Brightfield and epifluorescence imaging revealed characteristic monolayer formation on PDMS slabs, with cobblestone cell morphology, regularly spaced nuclei, and no observable differences between control and 3D printed replicas (**Figure 5a, d**). Consistent with these findings, cell viability measurements showed no significant differences for MCF10A cells (**Figure 5b**) or 4T1 cells (**Figure 5c**) between PDMS replicas from control and 3D printed surfaces coated with XTC-3D. Similarly, no differences were observed for MCF10A cells (**Figure 5e**) or 4T1 cells (**Figure 5f**) on PDMS replicas coated with Janchun epoxy. These findings demonstrate that PDMS replicas generated from epoxy-coated 3D printed molds support cell viability and monolayer formation comparable to control PDMS surfaces. By examining two distinct cell lines and two epoxy formulations (0.5 acetone mixtures with 1. XTC-3D epoxy and 2. Janchun epoxy), we confirmed the robustness of this approach for fabricating biocompatible PDMS substrates suitable for biological applications.

**Figure 5.**
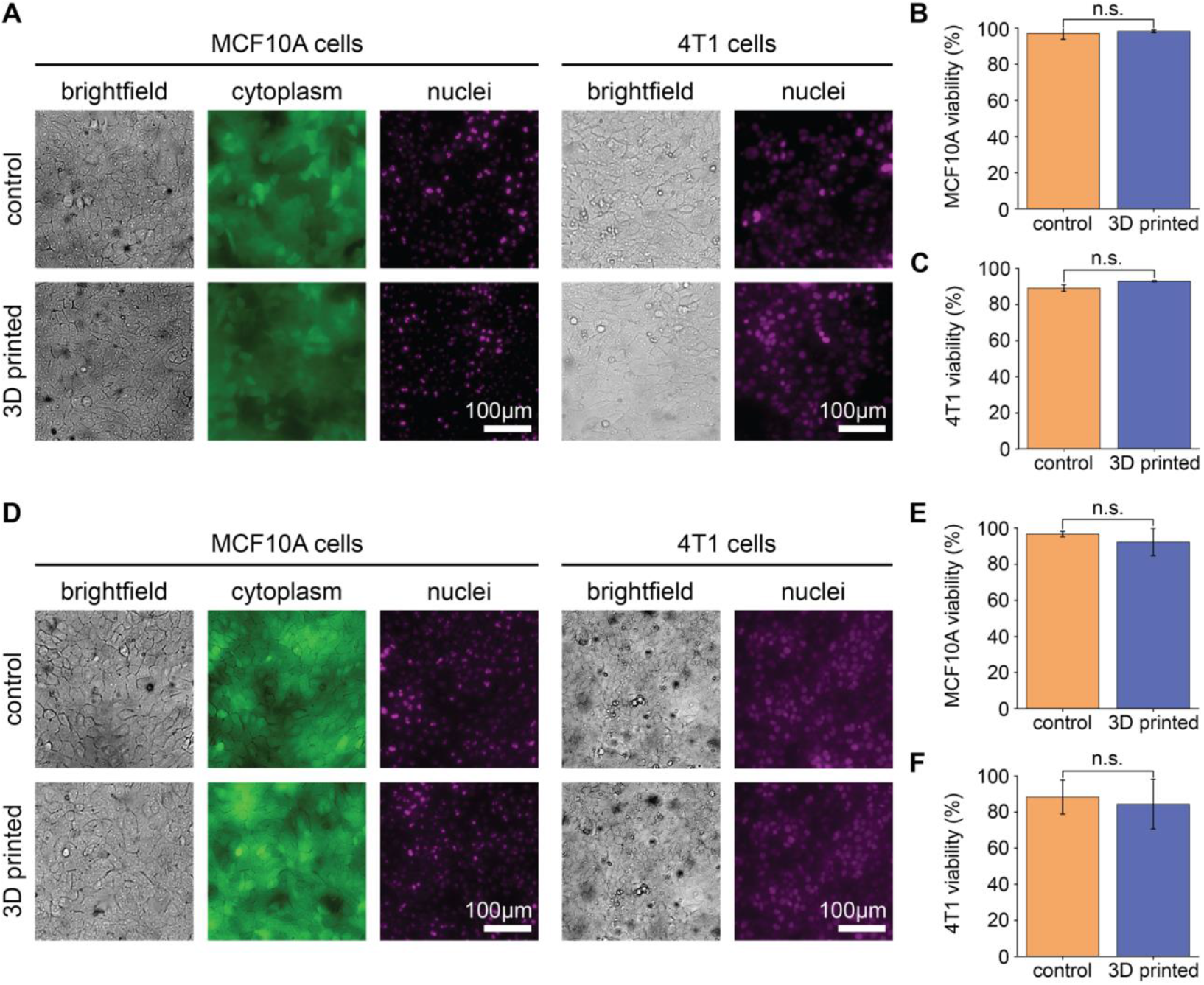
Biocompatibility of PDMS replicas fabricated using epoxy-coated 3D printed molds. Evaluation of MCF10A and 4T1 cell morphology and viability after 72 hours of culture on PDMS slabs cast from 3D printed molds coated with epoxy-acetone 0.5 mixtures, for XTC-3D epoxy **(A–C)** or Janchun epoxy **(D–F)**, compared to control PDMS cast from standard 10 cm polystyrene dishes. **(A, D)** Brightfield and epifluorescence images of MCF10A and 4T1 cells show comparable morphology and confluency between control and 3D printed mold conditions, supporting the biocompatibility of PDMS replicas fabricated from epoxy-coated molds. **(B–C, E–F)** Bar graphs quantifying viability of MCF10A cells **(B, E)** and 4T1 **(C, F)** cells cultured on control vs. 3D printed mold conditions. No significant differences (n.s.) were observed. Data represent mean ± standard deviation. Epifluorescence images in panels A and D show live MCF10A cells expressing cytoplasmic GFP (green) and nuclear RFP (magenta), and 4T1 cells expressing nuclear iRFP670 (magenta).

### Enabling 3D Printed Platforms for Organ-on-a-Chip Arrays and Live-Cell Imaging

With the biocompatibility of PDMS replicas from epoxy-coated 3D printed molds confirmed, along with our previous data demonstrating their suitability for fluidic applications and high clarity for optical imaging, we established an OoC array platform designed for live-cell imaging. As a proof-of-principle, we engineered blind-ended collagen cavities housed within fluidic channels (1 mm width × 1 mm height × 20 mm length) as a scaffold for 3D cell aggregate formation. We engineered distinct cell aggregates comprising 4T1 cells and MCF10A cells, mimicking breast tumor and breast tissue, respectively (**Figure 6a**). In addition to the generation of 3D cell aggregates, the simple fluidic channel geometry used here has wide ranging applications for organ-on-a-chip and fluidic cultures. For instance, similar channels have been used to engineer luminal tissue structures and vessels^24, 31–42^, expose matrix embedded cells to controlled fluid flow conditions^43–47^, and serve as spaces for circulating cells^48–50^ (**Figure S4**).^24, 31–36, 43–46, 48–50^ Accordingly, we established a user-friendly platform to assemble arrays of OoC devices on a custom 3D printed plate, enabling live-cell monitoring of fluidic cultures at scale (**Figure 6b-d**). We designed a 3D printed mold to fabricate a 4 × 5 array of channels with 1 mm × 1 mm × 20 mm dimensions, strategically spaced to facilitate PDMS cutting and handling (**Figure 6b**). To allow for maximum accessibility of the platform, we designed a workflow that supports gravity-driven fluid flow, eliminating the need for external fluidic control. To do this, we generated a 3D printed mold comprising a 4 × 5 array of cylindrical posts of 8 mm diameter and 4 mm height, with inter-post row spacing matched to the inter-channel row spacing (**Figure 6b**). The 3D channel mold produces an array of 20 PDMS channels, which can be cut into 4 devices comprising 5 fluidic channels each. Fittingly, the 3D post mold yields an array of PDMS wells that can be cut into 4 strips of 5 wells each, which can be placed on top of the PDMS channels at a desired end, serving as a media reservoir for gravity-driven flow (**Figure 6c**). As such, these 3D printed molds enable the facile generation of multi-format OoC PDMS devices, supporting user flexibility for applications in pump-driven or gravity-driven fluidic flow.

**Figure 6.**
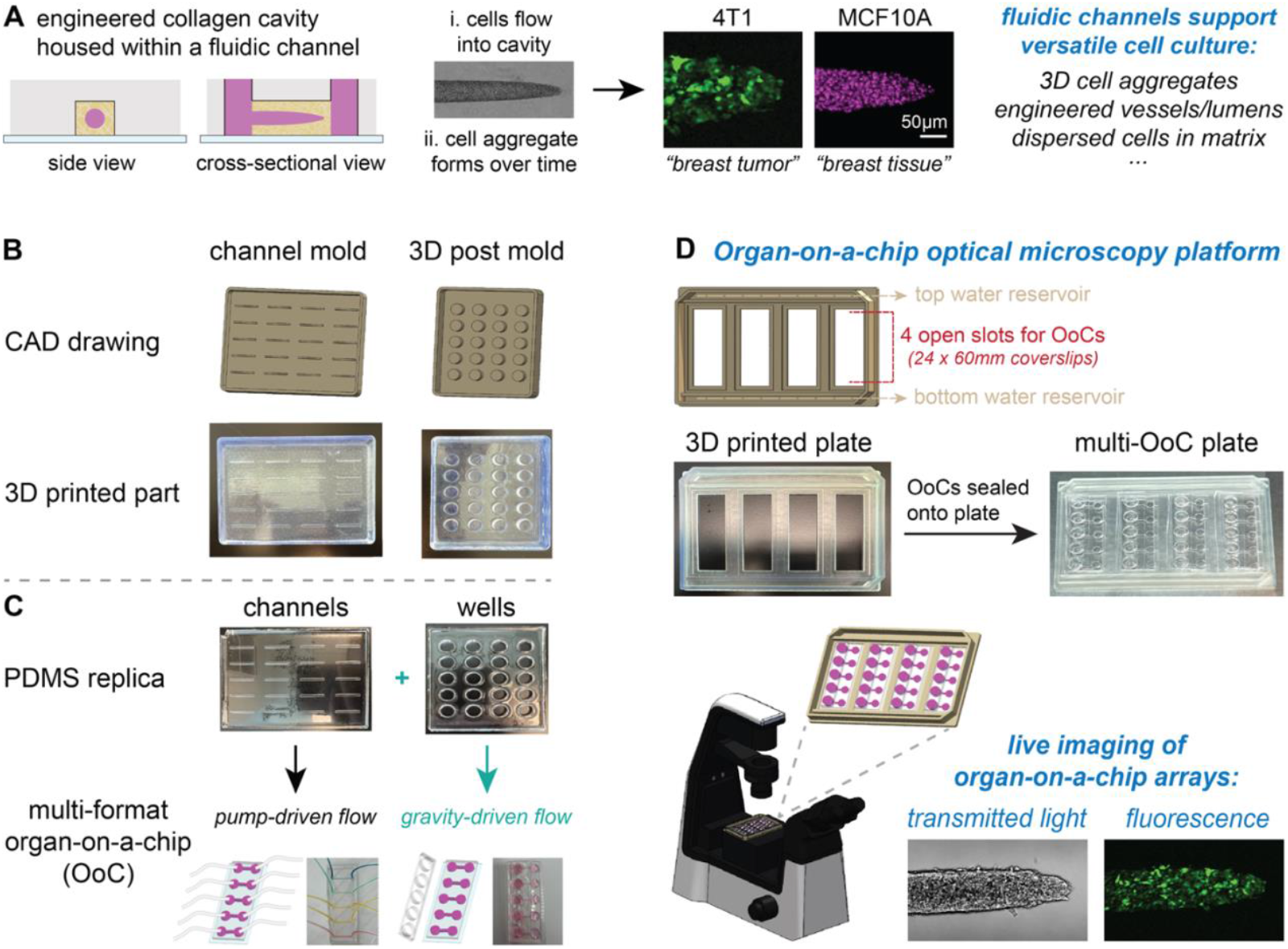
3D printed platform for streamlining OoC fabrication and live-cell imaging. **(A)** Workflow for seeding cells into an engineered collagen cavity housed within a fluidic channel. Side and cross-sectional schematics illustrate the cavity structure and flow of cells into the extracellular matrix, followed by aggregate formation over time; Z-projection images of confocal fluorescence for live 4T1 and MCF10A aggregates with endogenous nuclear expression of YFP (green) and RFP (magenta), respectively. This platform supports a range of biological applications, including 3D tissue models. **(B)** Illustrations of CAD drawings and 3D printed molds of channels and posts. **(C)** PDMS replicas produced from 3D printed molds enable modular integration of fluidic channel arrays into a multi-format organ-on-a-chip (OoC) platform compatible with both pump-driven and gravity-driven flow, in which arrays of fluidic channels are used for both formats; Channel connections to inlet and outlet tubing permit pump-driven flow, while PDMS wells from 3D posts placed on top of biopsy punched fluidic channels facilitate gravity-driven flow. **(D)** OoC optical microscopy platform with a custom 3D printed plate designed to hold four 24 × 60 mm coverslips and maintain humidity via top and bottom water reservoirs. Devices can be sealed onto the plate and imaged in parallel using transmitted light or fluorescence for live-cell monitoring of OoC arrays.

To facilitate downstream OoC applications, we designed a 3D printed plate that secures 4 glass coverslips in parallel and is compatible with live-cell monitoring using optical microscopy (**Figure 6d**). To ensure compatibility with existing laboratory equipment, the exterior dimensions of the 3D printed plate were informed by the dimensions of a standard well-plate (typically ~127 mm × ~85 mm), with platform wall height that accommodates a universal well plate lid. The plate bottom has four rectangular openings (24.5 mm × 60.5 mm) with a 1 mm interior overhang for placing OoC devices bonded to 24 mm × 60 mm glass coverslips. These coverslips may be secured to the plate using PDMS or vacuum grease as a sealant. Both methods are reversible, allowing for downstream applications and reuse of 3D printed imaging plates. Two reservoirs flank the regions above and below the plate’s rectangular windows, allowing for the addition of water to maintain a humidified chamber for live-cell imaging. This 3D printed platform allows for the simultaneous fabrication and monitoring of 20 OoCs, requiring only two 3D printed parts for pump-driven flow or three 3D printed parts for gravity-driven flow (**Figure 6d**). Notably, all 3D printed parts can be reused multiple times to fabricate and image fluidic devices, serving as a cost-effective tool.

## Discussion

Here, we established a facile approach for generating PDMS-based devices from 3D printed molds. Our method overcomes fabrication constraints associated with generating larger feature sizes in photolithography and directly using 3D printed parts for downstream applications in soft lithography. This approach utilizes epoxy alone or epoxy-acetone mixtures for coating 3D printed parts and is compatible with SLA and FDM printing. The application of epoxy coating, particularly at a 0.50 acetone ratio, significantly decreases surface imperfections and increases optical clarity of PDMS replicas from SLA printed molds. Fluidic channels with high structural integrity can be fabricated down to 400 µm cross-sectional dimensions, and 3D printed molds can be used to generate varied geometries for microarrays, with multitiered posts. Additionally, we optimized a two-step epoxy coating method for FDM printed molds. We demonstrated the robust casting of PDMS from 3D printed molds and irreversible bonding to glass to generate fluidic channels. Moreover, we confirmed the biocompatibility of PDMS replicas for cell culture applications, showing that cells can be directly cultured on top of PDMS replicas and housed within fluidic devices.

To facilitate OoC studies, we established a platform for the fabrication and assembly of millifluidic channels to serve as cell culture chambers for live cell monitoring. We include the design of 3D printed parts for both gravity-driven and pump-driven fluid flow. As a case study, we generated 3D cell aggregates in 3D matrix using normal and breast cancer cells within fluidic channels to demonstrate the suitability for OoCs. We designed a 3D printed plate to hold 4 glass coverslips in parallel that each house 5 OoCs, enabling 20 OoCs to be monitored at a time using live-cell imaging. With user care, the 3D printed parts can be reused multiple times to fabricate fluidic devices and conduct live-cell imaging, making this approach highly practical and cost-effective for diverse research laboratories. All 3D printed parts can be produced using basic 3D printers, and we provide open-source CAD drawings and STL files for facile assembly of OoC platforms. The complete set of CAD files for the 3D printed parts used in this study is publicly available via a GitHub repository (see **Data Availability**). By providing these resources, we significantly reduce barriers to entry, enabling researchers from various backgrounds to readily adopt and implement our OoC technology.

Our epoxy coating methods significantly expand the applicability of low-cost printing options, making this a particularly accessible method to a broad range of researchers, including those without prior microfabrication experience. The method is designed to be universal across different 3D printing platforms, including both SLA and FDM printers. This versatility allows for the fabrication of molds with varying sizes and complexities, serving as a highly customizable approach for the generation of PDMS-based devices. Accordingly, the primary limitations in PDMS device generation are imposed by the capabilities of the chosen 3D printer. For instance, SLA printers are known for their high resolution and precision, making them suitable for intricate designs, while FDM printers offer larger build volumes, accommodating bigger molds. However, it is important to acknowledge the limitations in resolution inherent to each printing technology. While SLA printers can achieve fine details, FDM printers may face challenges in replicating small features due to their layer height and nozzle diameter constraints. Despite these limitations, both printing methods have been successfully utilized in the fabrication of fluidic devices. For example, studies have demonstrated the use of FDM printing to create molds for PDMS microfluidic devices, highlighting its applicability in this field.^16, 17^

Looking forward, expanding this approach to incorporate a wider array of 3D printing technologies such as digital light processing (DLP), and liquid crystal display (LCD) printing could further diversify the design space and material compatibility of OoC platforms.^10, 51–53^ These advanced printing modalities enable the integration of soft and rigid materials, conductive elements, and multi-material gradients within a single device. Moreover, PDMS-based devices generated using 3D printed molds could serve as versatile cell culture platforms, integrating complementary technologies to enable precise spatial and temporal control over microenvironmental conditions to model physiological and pathological conditions in vitro.^54–56^ Notably, they offer opportunities to introduce mechanical actuation directly into the chip design. For instance, pressure-responsive chambers or strain-inducing geometries can be printed to apply tunable mechanical stresses that mimic physiological conditions such as peristalsis, blood flow, or tissue compression.^54, 57, 58^ Embedding pneumatic or hydraulic elements using PDMS and multi-layer printing could enable dynamic modulation of pressure, shear stress, and substrate stiffness. These mechanical cues are critical for simulating in vivo microenvironments, particularly in studies of cancer progression, vascular remodeling, or organ development.^25, 59^ Integrating such mechanical forces into 3D printed OoC systems opens new avenues for exploring mechanoresponsive signaling pathways and drug responses under more physiologically relevant conditions.

## Conclusions

This study highlights the potential of epoxy-coated 3D printed molds for the fabrication of PDMS-based devices, with applications in fluidics and OoC devices. By addressing long-standing challenges with casting PDMS from SLA and FDM printed molds, including surface quality, feature fidelity, optical clarity, and biocompatibility, our method establishes a robust framework for advancing the design and production of customizable and cost-effective PDMS-based devices. Our accessible and tunable platform for live imaging of OoC systems represents a significant step toward enabling scalable, high-throughput research in tissue engineering, drug development, and personalized medicine.^25^

## Author contributions

**R.J. Abbed:** Conceptualization, Data curation, Formal Analysis, Visualization, Investigation, Methodology, Writing – original draft, Writing – review & editing. **E.I. Quiñones Cruz:** Formal Analysis, Visualization, Investigation, Writing – original draft, Writing – review & editing. **S.E. Leggett:** Conceptualization, Resources, Project administration, Funding acquisition, Supervision, Writing – review & editing.

## Data availability

The CAD drawings and STL files used to fabricate the 3D printed components in this study are available in the v1.0 release of our GitHub repository. All files required to replicate the device designs are included. Please cite this release as: Abbed, R.J., Quiñones Cruz, E., Leggett, S.E. *3D Printed Molds for Organ-on-a-Chip and Fluidics: PDMS-Based Rapid and Accessible Prototyping*. GitHub. Release v1.0., 2025. https://github.com/leggettlab/3D-Printed-Molds-for-Organ-on-a-Chip-and-Fluidics-PDMS-Based-Rapid-and-Accessible-Prototyping.

## Acknowledgements

We thank I.Y. Wong for insightful comments, as well as D.A. Haber for the MCF10A cell line expressing cytoplasmic GFP and RFP H2B, and C.M. Nelson for the 4T1 iRFP670 cell line. This work was supported by faculty start-up funds to S.E.L. from the University of Illinois Urbana-Champaign and the Cancer Center at Illinois.

## References

1. A. V. Nielsen, M. J. Beauchamp, G. P. Nordin and A. T. Woolley, Annu Rev Anal Chem (Palo Alto Calif), 2020, 13, 45–65.

2. A. L. Beckwith, L.F. Velásquez-García and J. T. Borenstein, Adv Healthc Mater, 2019, 8, e1900289.

3. C. S. Carrell, C. P. McCord, R. M. Wydallis and C. S. Henry, Anal Chim Acta, 2020, 1124, 78–84.

4. M. Mellody, Y. Nakagawa, R. James and D. Di Carlo, Lab Chip, 2025, 25, 1565–1574.

5. R. Hernández Vera, P. O’Callaghan, N. Fatsis-Kavalopoulos and J. Kreuger, Sci Rep, 2019, 9, 11321.

6. B. V. Dang, A. Hassanzadeh-Barforoushi, M. S. Syed, D. Yang, S. J. Kim, R. A. Taylor, G. J. Liu, G. Liu and T. Barber, ACS Sens, 2019, 4, 2181–2189.

7. A. K. Au, W. Huynh, L. F. Horowitz and A. Folch, Angewandte Chemie, 2016, 55, 3862–3881.

8. B. J. O’Grady, M. D. Geuy, H. Kim, K. M. Balotin, E. R. Allchin, D. C. Florian, N. N. Bute, T. E. Scott, G. B. Lowen, C. M. Fricker, M. L. Fitzgerald, S. A. Guelcher, J. P. Wikswo, L. M. Bellan and E. S. Lippmann, Lab Chip, 2021, 21, 4814–4822.

9. A. Ahmadianyazdi, I. J. Miller and A. Folch, Lab Chip, 2023, 23, 4019–4032.

10. H. Shafique, V. Karamzadeh, G. Kim, M. L. Shen, Y. Morocz, A. Sohrabi-Kashani and D. Juncker, Lab Chip, 2024, 24, 2774–2790.

11. M. Ramasamy, B. Ho, C.-M. Phan, N. Qin, L. C. Ren and L. Jones, Journal of Micromechanics and Microengineering, 2023, 33.

12. N. H. Chan, Y. Chen, Y. Shu, Y. Chen, Q. Tian and H. Wu, 2015, 19, 9–18.

13. G. Comina, A. Suska and D. Filippini, Lab Chip, 2014, 14, 424–430.

14. D. H. Han, U. Oh and J. K. Park, ACS Omega, 2023, 8, 19128–19136.

15. K. Kamei, Y. Mashimo, Y. Koyama, C. Fockenberg, M. Nakashima, M. Nakajima, J. Li and Y. Chen, Biomed Microdevices, 2015, 17, 36.

16. O. Riester, S. Laufer and H. P. Deigner, J Nanobiotechnology, 2022, 20, 540.

17. R. F. Quero, G. Domingos da Silveira, J. A. Fracassi da Silva and D. P. Jesus, Lab Chip, 2021, 21, 3715–3729.

18. I. Miranda, A. Souza, P. Sousa, J. Ribeiro, E. M. S. Castanheira, R. Lima and G. Minas, J Funct Biomater, 2021, 13.

19. M. de Almeida Monteiro Melo Ferraz, J. B. Nagashima, B. Venzac, S. Le Gac and N. Songsasen, Sci Rep, 2020, 10, 994.

20. D. W. Simmons, D. R. Schuftan, G. Ramahdita and N. Huebsch, ACS Appl Mater Interfaces, 2023, 15, 25313–25323.

21. G. van der Borg, H. Warner, M. Ioannidis, G. van den Bogaart and W. H. Roos, Polymers (Basel), 2023, 15.

22. M. Villegas, Z. Cetinic, A. Shakeri and T. F. Didar, Anal Chim Acta, 2018, 1000, 248–255.

23. R. Salazar, F. Pizarro, D. Vasquez and E. Rajo-Iglesias, Additive Manufacturing, 2022, 51, 102593.

24. S. E. Leggett, M. C. Brennan, S. Martinez, J. Tien and C. M. Nelson, Cell Mol Bioeng, 2024, 17, 7–24.

25. B. C. H. Cheung, R. J. Abbed, M. Wu and S. E. Leggett, Annu Rev Biomed Eng, 2024, 26, 93–118.

26. H. B. Musgrove, M. A. Catterton and R. R. Pompano, Anal Chim Acta, 2022, 1209, 339842.

27. H. B. Musgrove, S. R. Cook and R. R. Pompano, ACS Appl Bio Mater, 2023, 6, 3079–3083.

28. J. Debnath, S. K. Muthuswamy and J. S. Brugge, Methods, 2003, 30, 256–268.

29. S. Javaid, J. Zhang, E. Anderssen, J. C. Black, B. S. Wittner, K. Tajima, D. T. Ting, G. A. Smolen, M. Zubrowski, R. Desai, S. Maheswaran, S. Ramaswamy, J. R. Whetstine and D. A. Haber, Cell Rep, 2013, 5, 1679–1689.

30. A. S. Piotrowski-Daspit, A. K. Simi, M. F. Pang, J. Tien and C. M. Nelson, Methods Mol Biol, 2017, 1501, 245–257.

31. J. Cacheux, T. Nakajima, D. Alcaide, T. Sano, K. Doi, A. Bancaud and Y. T. Matsunaga, STAR Protoc, 2024, 5, 102950.

32. Y. W. Dance, M. C. Obenreder, A. J. Seibel, T. Meshulam, J. W. Ogony, N. Lahiri, L. Pacheco-Spann, D. C. Radisky, M. D. Layne, S. R. Farmer, C. M. Nelson and J. Tien, Cell Mol Bioeng, 2023, 16, 23–39.

33. J. M. Ayuso, M. M. Gong, M. C. Skala, P. M. Harari and D. J. Beebe, Adv Healthc Mater, 2020, 9, e1900925.

34. J. M. Ayuso, S. Rehman, M. Farooqui, M. Virumbrales-Muñoz, V. Setaluri, M. C. Skala and D. J. Beebe, Int J Mol Sci, 2020, 21.

35. K.M. Lugo-Cintrón, J. M. Ayuso, B. R. White, P. M. Harari, S. M. Ponik, D. J. Beebe, M. M. Gong and M. Virumbrales-Muñoz, Lab Chip, 2020, 20, 1586–1600.

36. J. Tien, U. Ghani, Y. W. Dance, A. J. Seibel, M. Karakan, K. L. Ekinci and C. M. Nelson, iScience, 2020, 23, 101673.

37. E. Delannoy, G. Tellier, J. Cholet, A. M. Leroy, A. Treizebré and F. Soncin, Biomedicines, 2022, 10.

38. T. J. Kwak and E. Lee, Sci Rep, 2020, 10, 20142.

39. T. Mathur, K. A. Singh, N. K. R Pandian, S. H. Tsai, T. W. Hein, A. K. Gaharwar, J. M. Flanagan and A. Jain, Lab Chip, 2019, 19, 2500–2511.

40. C. G. M. van Dijk, M. M. Brandt, N. Poulis, J. Anten, M. van der Moolen, L. Kramer, E. F. G. A. Homburg, L. Louzao-Martinez, J. Pei, M. M. Krebber, B. W. M. van Balkom, P. de Graaf, D. J. Duncker, M. C. Verhaar, R. Luttge and C. Cheng, Lab Chip, 2020, 20, 1827–1844.

41. M. Virumbrales-Muñoz, J. Chen, J. Ayuso, M. Lee, E. J. Abel and D. J. Beebe, Lab Chip, 2020, 20, 4420–4432.

42. R. G. Mannino, D. R. Myers, B. Ahn, Y. Wang, Margo Rollins, H. Gole, A. S. Lin, R. E. Guldberg, D. P. Giddens, L. H. Timmins and W. A. Lam, Sci Rep, 2015, 5, 12401.

43. M. Pandey, Y. J. Suh, M. Kim, H. J. Davis, J. E. Segall and M. Wu, Phys Biol, 2024, 21.

44. G. Saorin, I. Caligiuri and F. Rizzolio, Semin Cell Dev Biol, 2023, 144, 41–54.

45. S. E. Leggett, M. Patel, T. M. Valentin, L. Gamboa, A. S. Khoo, E. K. Williams, C. Franck and I. Y. Wong, Proc Natl Acad Sci U S A, 2020, 117, 5655–5663.

46. Y. L. Huang, Y. Ma, C. Wu, C. Shiau, J. E. Segall and M. Wu, Sci Rep, 2020, 10, 9648.

47. A. Markoski, I. Y. Wong and J. T. Borenstein, Front Med Technol, 2021, 3, 646441.

48. M. Wang, T. Zhu, C. Liu, L. Jin, P. Fei and B. Zhang, Biomed Pharmacother, 2022, 154, 113567.

49. B. P. Rickard, C. Conrad, A. J. Sorrin, M. K. Ruhi, J. C. Reader, S. A. Huang, W. Franco, G. Scarcelli, W. J. Polacheck, D. M. Roque, M. G. Del Carmen, H. C. Huang, U. Demirci and I. Rizvi, Cancers (Basel), 2021, 13.

50. I. Rizvi, U. A. Gurkan, S. Tasoglu, N. Alagic, J. P. Celli, L. B. Mensah, Z. Mai, U. Demirci and T. Hasan, Proc Natl Acad Sci U S A, 2013, 110, E1974–1983.

51. Z. Luo, H. Zhang, R. Chen, H. Li, F. Cheng, L. Zhang, J. Liu, T. Kong, Y. Zhang and H. Wang, Microsyst Nanoeng, 2023, 9, 103.

52. Y. T. Kim, A. Ahmadianyazdi and A. Folch, Nat Protoc, 2023, 18, 1243–1259.

53. P. Prabhakar, R. K. Sen, N. Dwivedi, R. Khan, P. R. Solanki, A. K. Srivastava and C. Dhand, Frontiers in Nanotechnology, 2021, 3.

54. A. H. Szmelter, G. Venturini, R. J. Abbed, M. O. Acheampong and D. T. Eddington, Lab Chip, 2023, 23, 793–802.

55. A. Szmelter, J. Jacob and D. T. Eddington, Anal Chem, 2021, 93, 2570–2577.

56. T. M. Valentin, S. E. Leggett, P. Y. Chen, J. K. Sodhi, L. H. Stephens, H. D. McClintock, J. Y. Sim and I. Y. Wong, Lab Chip, 2017, 17, 3474–3488.

57. C. Rein, M. Toner and D. Sevenler, Sci Rep, 2023, 13, 1232.

58. B. R. Mutlu, T. Dubash, C. Dietsche, A. Mishra, A. Ozbey, K. Keim, J. F. Edd, D. A. Haber, S. Maheswaran and M. Toner, Lab on a Chip, 2020, 20, 1612–1620.

59. C. L. Thompson, S. Fu, M. M. Knight and S. D. Thorpe, Front Bioeng Biotechnol, 2020, 8, 602646.

