## Supplemental Figures 1-4 for "3D Printed Molds for Organ-on-a-Chip and Fluidics: PDMS-Based Rapid and Accessible Prototyping"

**Rana J. Abbed[1], Edwin I. Quiñones Cruz[1], and Susan E. Leggett[1,2*]**

[1]Department of Bioengineering, University of Illinois Urbana-Champaign, 1406 W. Green St., Urbana, IL, USA,

[2]Cancer Center at Illinois, University of Illinois Urbana-Champaign, 405 N. Mathews Ave., Urbana, IL, USA

*** Corresponding author:**
S.E. Leggett
Department of Bioengineering
Cancer Center at Illinois
University of Illinois Urbana-Champaign,
2240 Everitt Laboratory
1406 W. Green St.
Urbana, IL, 61801 USA


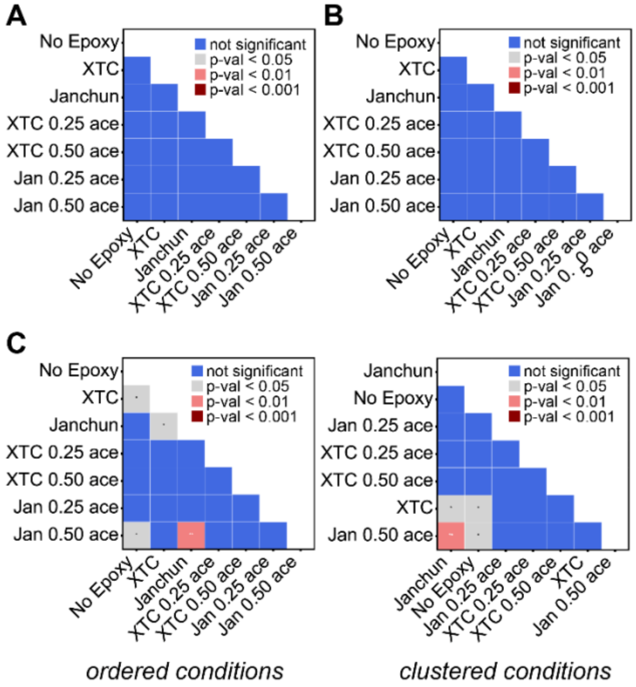


**Supplemental Figure 1. Statistical comparison of dimensional fidelity for PDMS replicas fabricated from SLA printed molds. (A-C)** Pairwise significance heatmaps compare statistical differences in channel dimensions across different epoxy coating conditions for **(A)** length, **(B)** width, and **(C)** height. **(A-B)** Heatmaps show no significant differences for channel length and width dimensions; conditions are shown in the order they are displayed for the corresponding data plot in Fig. 2 D-E. **(C)** Heatmaps comparing channel height dimensions using two layouts: ordered conditions (left) and clustered conditions (right). The ordered layout displays conditions shown in the order they appear in the corresponding data plot in Fig. 2F, while the clustered layout groups conditions for ease of visualization of statistically different comparisons. Block colors and asterisks indicate the level of statistical significance: blue = not significant, gray/* = p-value < 0.05, pink/** = p-value < 0.01, and red/*** = p-value < 0.001. Abbreviated terms represent XTC-3D (XTC), Janchun (Jan), and acetone (ace).

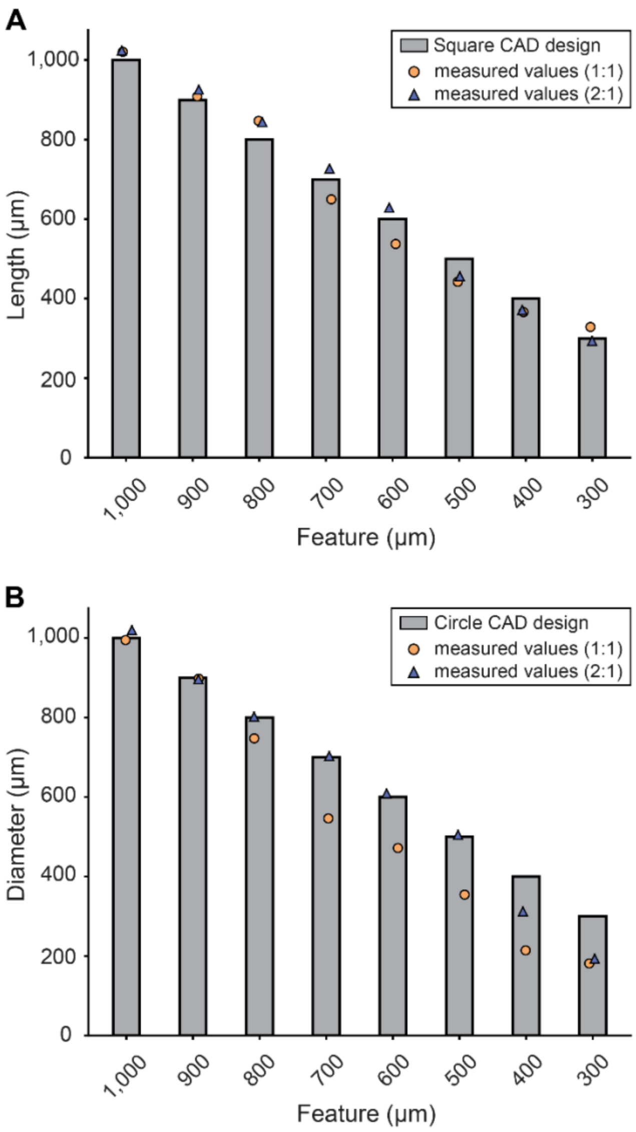


**Supplemental Figure 2. Fabrication fidelity of square and circular geometries from SLA printed molds. (A-B)** Bar graphs comparing nominal CAD dimensions (gray bars) to measured values for square features **(A)** and circular features **(B)** with 1:1 (orange circles) and 2:1 (blue triangles) aspect ratios of height to side length and diameter, respectively. Circular features exhibit greater deviation at smaller scales, particularly at a 1:1 aspect ratio.

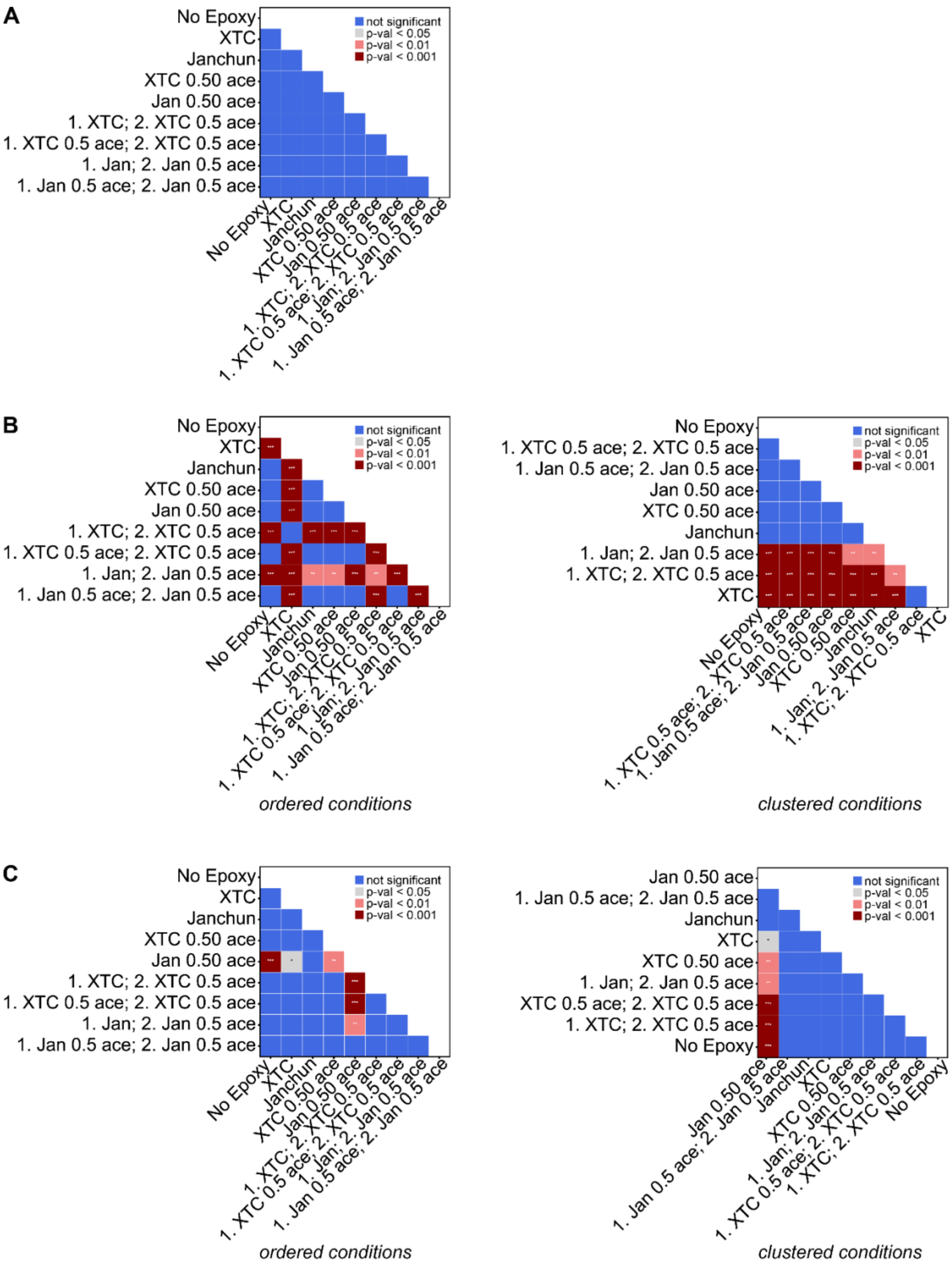


**Supplemental Figure 3. Statistical comparison of dimensional fidelity for PDMS replicas fabricated from FDM printed molds. (A- C)** Pairwise significance heatmaps compare statistical differences in channel dimensions across different epoxy coating conditions for **(A)** length, **(B)** width, and **(C)** height. Heatmaps comparing channel dimensions using two layouts: ordered conditions (left) and clustered conditions (right). The ordered layout displays conditions shown in the order they appear in the corresponding data plot in Fig. 4D-F, while the clustered layout groups conditions for ease of visualization of statistically different comparisons. Block colors and asterisks indicate the level of statistical significance: blue = not significant, gray/* = p-value < 0.05, pink/** = p-value < 0.01, and red/*** = p-value < 0.001. Abbreviated terms represent XTC-3D (XTC), Janchun (Jan), and acetone (ace).

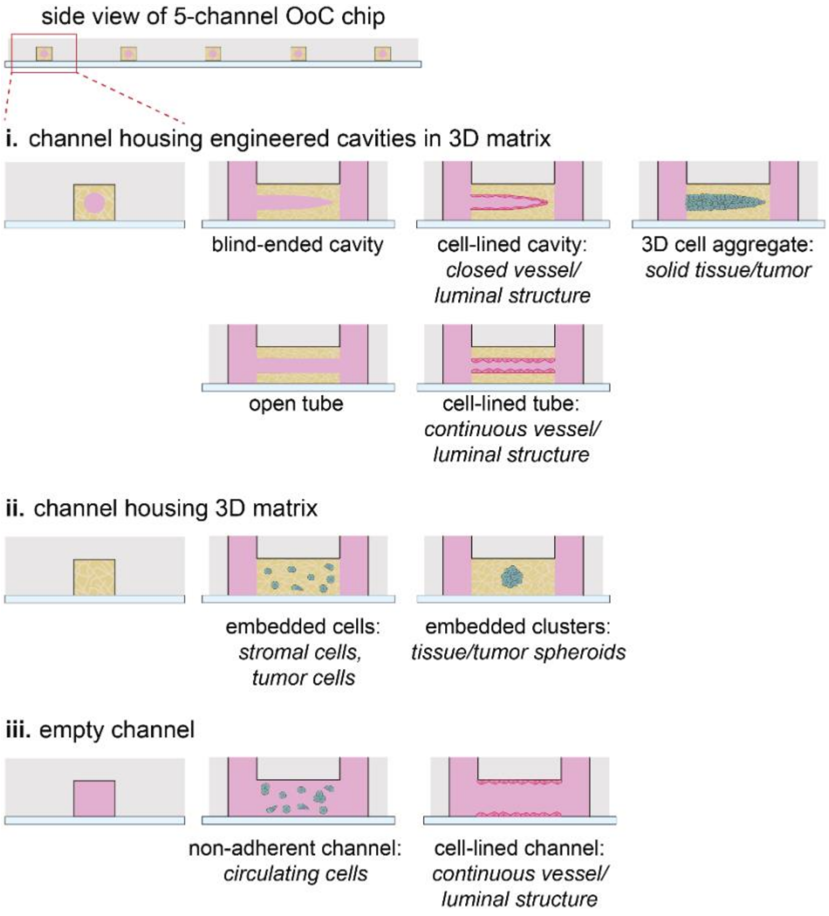


**Supplemental Figure 4. Organ-on-a-chip configurations and applications enabled by our accessible 3D printing platform for fabrication of PDMS-based fluidic devices.** Side view of a 5-channel OoC chip and **(i–iii)** magnified side and longitudinal cross-sectional views illustrating diverse tissue engineering formats supported by our platform. **(i)** Channels housing engineered cavities within a 3D matrix, demonstrating strategies to create perfusable 3D culture environments. These configurations support a range of tissue structures, including blind-ended cavities for forming cell-lined compartments resembling closed vessels or luminal structures, solid 3D cell aggregates representing tissues or tumors, open tubes, and continuous cell-lined tubes mimicking vascular networks. **(ii)** Channels filled entirely with 3D matrix to support embedded cells such as dispersed single cells, multicellular clusters, or spheroids. **(iii)** Hollow fluidic channels without 3D matrix for applications such as circulating cell studies in non-adherent channels or formation of continuous luminal structures in cell-lined channels, mimicking vascular systems.
